# Necroptosis inhibition counteracts neurodegeneration, memory decline and key hallmarks of aging, promoting brain rejuvenation

**DOI:** 10.1101/2021.11.10.468052

**Authors:** Macarena S. Arrázola, Matías Lira, Gabriel Quiroz, Felipe Véliz-Valverde, Somya Iqbal, Samantha L Eaton, Rachel A Kline, Douglas J Lamont, Hernán Huerta, Gonzalo Ureta, Sebastián Bernales, J César Cárdenas, Waldo Cerpa, Thomas M. Wishart, Felipe A. Court

## Abstract

Age is the main risk factor for the development of neurodegenerative diseases. In the aged brain, axonal degeneration is an early pathological event, preceding neuronal dysfunction, and cognitive disabilities in humans, primates, rodents, and invertebrates. Necroptosis mediates degeneration of injured axons, but whether necroptosis triggers neurodegeneration and cognitive impairment along aging is unknown. Here we show that the loss of the necroptotic effector *Mlkl* was sufficient to delay age-associated axonal degeneration and neuroinflammation, protecting against decreased synaptic transmission and memory decline in aged mice. Moreover, short-term pharmacologic inhibition of necroptosis in aged mice reverted structural and functional hippocampal impairment, both at the electrophysiological and behavioral level. Finally, a quantitative proteomic analysis revealed that necroptosis inhibition leads to an overall improvement of the aged hippocampal proteome, including a subclass of molecular biofunctions associated with brain rejuvenation, such as long-term potentiation and synaptic plasticity. Our results demonstrate that necroptosis contributes to the age-dependent brain degeneration, disturbing hippocampal neuronal connectivity, and cognitive function. Therefore, necroptosis inhibition constitutes a potential geroprotective strategy to treat age-related disabilities associated with memory impairment and cognitive decline.

## 1. INTRODUCTION

The current rise in human life expectancy is not precisely accompanied by an equivalent increase in healthspan (Aburto et al. 2020), the functional and disease-free period of life in the elderly (Hansen & Kennedy 2016). The impact of age on brain function is unquestionable, being the main risk factor for the development of neurodegenerative diseases and cognitive disabilities (Duan et al. 2020; Agüero-Torres et al. 2002; Wyss-Coray 2016).

As a fundamental structure for human cognition, the hippocampus is particularly vulnerable to the deleterious effects of aging (O’Shea et al. 2016; Wimmer et al. 2012). Microstructural changes at the synaptic level are correlated with learning and memory impairment during aging. Decreased number of axospinous synapses have been described in the aged hippocampus (Geinisman et al. 1992). The reduced number of synaptic contacts is correlated with a decreased presynaptic fiber potential due to a reduction of axons (Barnes & McNaughton 1980), contributing to the impaired synaptic plasticity and cognitive deficits evidenced in aged organisms (Rosenzweig & Barnes 2003). In addition, white matter abnormalities and axonal degeneration (AxD) have been identified in aged brains of diverse species (Peters et al. 2000; Stahon et al. 2016), particularly in the hippocampus, and correlated with impaired memory performance in humans (Radhakrishnan et al. 2020). Due to the importance of axonal integrity on hippocampal function and progression of cognitive decline in the elderly (Marner et al. 2003), it is imperative to determine the mechanism of AxD during aging. We have shown that necroptosis is involved in mechanical and chemical-induced AxD (Arrázola et al. 2019; Hernández et al. 2018). Recently, in a model of neuronal inflammation it has been demonstrated that necroptosis activates Sarm1, a central executioner of pathological AxD (Gerdts et al. 2013; Osterloh et al. 2012; Ko et al. 2020). Necroptosis is an alternative form of programmed cell death triggered by the tumor necrosis factor under caspase-8 inhibitory conditions and characterized by a necrotic-like and pro-inflammatory response (Holler et al. 2000; Seo et al. 2021). Receptor-interacting kinase 1 (RIPK1) recruits and phosphorylates RIPK3, which in turn phosphorylates the mixed lineage kinase domain-like protein (MLKL). MLKL oligomerizes and translocate to the plasma membrane, disrupting membrane integrity followed by the release of cellular components, an exacerbated inflammatory response, and cell death (Samson et al. 2020).

Age-associated increase in low-grade sterile inflammation contributes to the progression of age-associated diseases (Kennedy et al. 2014; Franceschi & Campisi 2014; López-Otín et al. 2013). Recent studies have depicted the importance of necroptosis in the aging of the mouse male reproductive system (Li et al. 2017) and the epididymal white adipose tissue (Deepa et al. 2018). In the context of brain aging, several age-related neurodegenerative conditions with prominent AxD and neuroinflammation as common features have shown increased necroptosis activation in the brain, associated with functional impairment (Re et al. 2014; Ito et al. 2016; Iannielli et al. 2018; Oñate et al. 2020; Caccamo et al. 2017; Salvadores et al. 2022). Necroptosis activation has been recently described in the cortex and hippocampus of aged mice, and correlated to age-related neuroinflammation (Thadathil et al. 2021), and in the basolateral amygdala in a senescence-accelerated mouse model of aging, and associated to reduced social interaction (Zhang et al. 2022). However, the involvement of necroptosis in the progression of normal brain aging and its impact on the age-associated memory decline remain unexplored.

Here, we investigated the role of necroptosis in the progression of neurodegeneration in the hippocampus and its impact in brain function along aging. Necroptosis increased in the hippocampus of aged mice and accompanied with evident features of AxD. Loss of *Mlkl* was sufficient to delay age-related AxD and neuroinflammation in the hippocampus, a youthful phenotype also displayed at the synaptic and functional level. Restored synaptic transmission and facilitation were accompanied by improved learning and memory in aged mice deficient for *Mlkl*. Short-term inhibition of RIPK3 in aged mice demonstrated to be extraordinarily effective on reverting neurodegeneration and hippocampus-dependent functional impairment. Finally, an unbiased quantitative proteomic analysis demonstrated that necroptosis inhibition improved the aged hippocampal proteomic profile, restoring the levels of key protein pathways associated with brain aging. Our study reveals that necroptosis contributes to the age-associated deterioration of axons, affecting hippocampal neuronal connectivity and cognitive functions, such as learning and memory performance, of aged mice. Therefore, necroptosis inhibition constitutes a potential strategy for the development of therapeutic tools to treat age-related brain disabilities and brain rejuvenation.

## 2. RESULTS

### 2.1. Necroptosis activation in the hippocampus of the aged mice

To establish the progression of hippocampal degeneration through aging we used adult (3-6 months), old (12-15 months) and aged (more than 20 months) mice. Neurodegeneration, evaluated by Fluoro Jade C (FJC) staining significantly increases in the hippocampus of aged mice (**Fig S1**). FJC staining also increased along aging in other brain regions, including the striatum, cerebellum and spinal cord (**Fig S1**). In order to evaluate age-associated AxD in the hippocampus, we studied the expression of two phosphorylated forms of NFs: highly phosphorylated NFs which are predominant in stable axons, while increased non-phosphorylated NF (non-pNF) immunoreactivity is associated with axonal damage (Nadeem et al. 2016; Petzold 2005). Axonal integrity was analyzed by calculating the Axonal Integrity Index (**Fig S2**) from pan-axonal NF-stained axons (stable axonal NFs) in the hippocampus (**Fig 1a**). The integrity and percentage of NF+ axons decreased in old and aged hippocampus compared with adult mice (**Fig 1a,c,d**). Almost undetectable immunoreactivity of non-pNF was observed in the hippocampus of adult mice, while a progressive increase was evidenced during aging in different subfields of the hippocampus (**Fig 1b, e**).

**Fig 1.**
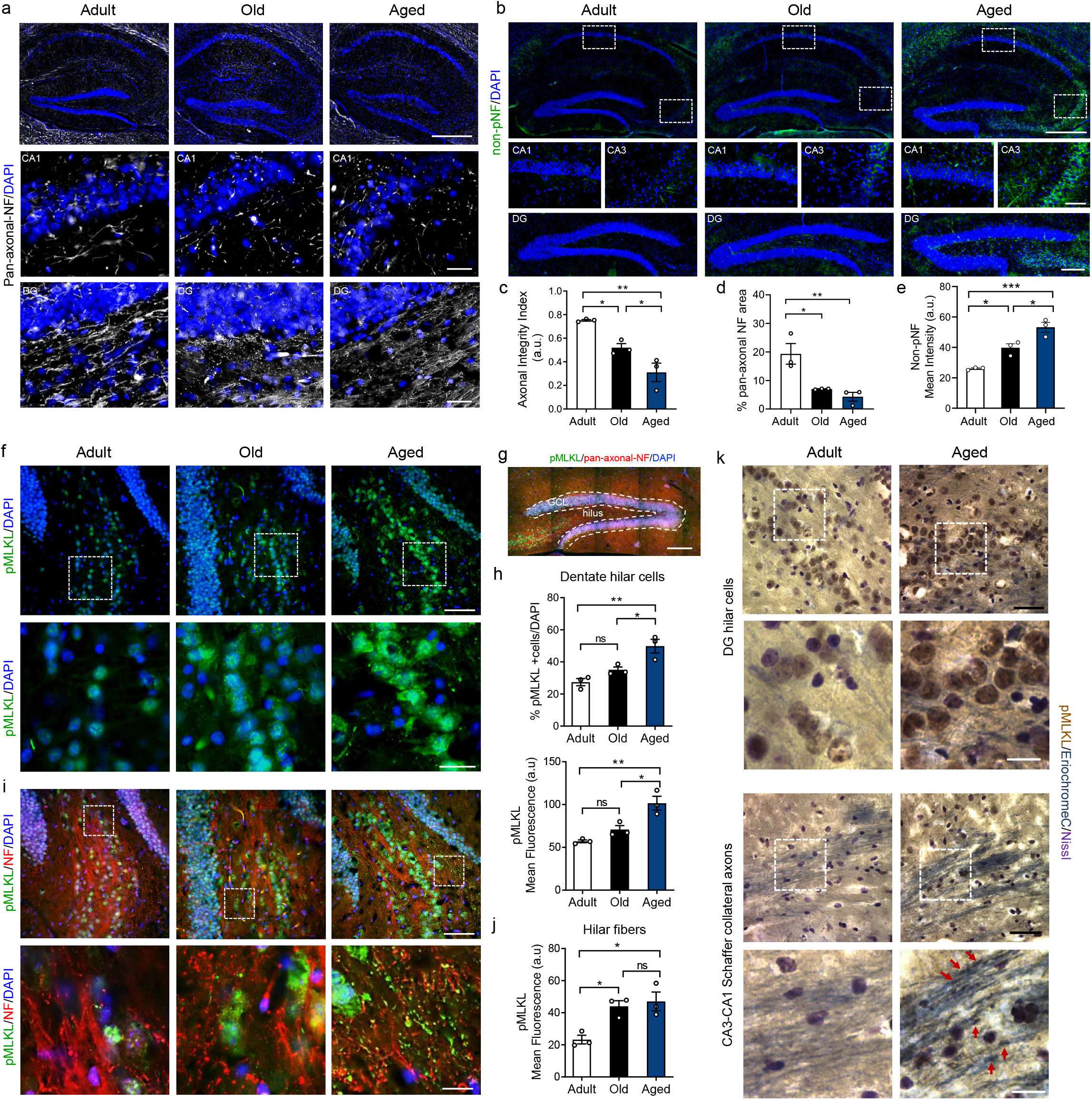
Progression of axonal degeneration coupled with necroptosis activation in the hippocampus during aging. Hippocampal sections of adult, old and aged mice stained with **(a)** pan-axonal NF (bars: 500 μm, hippocampus; 10 μm, crop), and **(b)** non-phosphorylated neurofilament (non-pNF) (bars: 500 μm, hippocampus; 200 μm, DG; 100 μm, CA1/CA3). DAPI, nuclei (blue). **(c)** Axonal integrity index and percentage of axonal area in the hippocampus **(d)**. **(e)** Non-pNF mean intensity in the hilus along aging. **(f)** MLKL phosphorylation (pMLKL, green; bars: 25 μm; 10 μm for magnification). **(g)** Mosaic reconstruction of aged hippocampal DG showing pMLKL in the hilus and granular cell layer (GCL) (green, bar: 100 μm). **(h)** Percentage of pMLKL positive hilar cells normalized to the total cell number in the hilar area (upper graph). pMLKL mean intensity (lower graph). **(i)** Axonal pMLKL detected in NF-positive axons, dotted boxes: magnified areas (bars: 25 μm and 5 μm, inset). **(j)** pMLKL mean intensity in hilar fibers along aging. **(k)** Immunohistochemistry against pMLKL (brown) and Eriochrome-C staining (myelinated axons, blue) (bar: 40 μm). Red dotted boxes, image magnification (bar: 15 μm). pMLKL in DG axons and in Schaffer collateral projections (red arrowheads). Increased pMLKL in neuronal somas (dark brown/+Nissl, violet). Values, mean ± SEM, are the result of the analysis of n=3-4 mice per group. One-way ANOVA with Tukey analysis for multiple comparisons, *p:<0.05; **p<0.01; ***p<0.005; ****p<0.001.

We recently demonstrated that mechanical and chemical-induced axonal degeneration is regulated by necroptosis (Arrázola et al. 2019). To determine whether necroptosis is involved in age-associated AxD, we assessed necroptosis activation by the expression of the phosphorylated form of MLKL (pMLKL) in the different hippocampal subfields, including the hilar axons of the DG and the CA3-CA1 projecting Schaffer collateral axons (**Fig 1f-k**). The number of pMLKL positive hilar cells increased in aged mice, as did pMLKL mean intensity (**Fig 1f, h**). Necroptosis activation was also evidenced by an age-dependent translocation of pMLKL from the nucleus to the cytoplasm (**Fig 1f**, magnified image), as previously described (Yoon et al. 2016). Axonal pMLKL staining was evaluated in pan-axonal-NF positive hilar fibers of the DG (**Fig 1i**). Interestingly, pMLKL mean intensity increased earlier in DG axons compared with dentate hilar somas throughout aging, reaching significant differences in the old mice group (**Fig 1h**). Increased pMLKL signal was also observed in other brain regions with defined axonal subfields, as axonal tracts in the striatum, the cerebellar white matter and the ventral horn of the spinal cord, which also showed progressive AxD along aging (**Fig S3**). The increase in pMLKL levels in the hilus was also accompanied by changes in the pattern of pMLKL signal, from almost non-detected in adult mice axons, diffuse in the old group, to finally become punctuated in axons of aged mice (**Fig 1i**, magnified image). These pMLKL aggregates have been associated with MLKL oligomerization and its translocation to the plasma membrane, two key steps for necroptosis execution (Samson et al. 2020). Same pMLKL pattern was observed by immunohistochemistry against pMLKL in DG hilar cells and in the Schaffer collateral axons of CA3-CA1 circuit of aged mice (**Fig 1k**). These results indicate that necroptosis is activated during aging in the hippocampus, a brain region that is considered one of the most vulnerable to the detrimental effects of aging, affecting learning and memory (Spiegel et al. 2013).

### 2.2. Age-induced axonal degeneration and neuroinflammation is reduced in MLKL knockout mice

Due to the indispensable role of MLKL in executing necroptosis, we evaluated whether age-dependent AxD in the hippocampus was modified in aged *Mlkl-knockout* mice (*Mlkl-KO*) (Wu et al. 2013). Interestingly, AxD in pan-axonal NF-stained axons was significantly reduced in aged *Mlkl-KO* mice at levels comparable with adult mice (**Fig 2a, b**). Moreover, non-pNF degenerated axons profusely present in aged WT hippocampus, were almost undetected in aged *Mlkl-KO* mice, at levels equivalent to younger WT mice (**Fig 2a, c**). As expected, we observed age-dependent AxD coupled with neuroinflammation (Hwang et al. 2018) (**Fig S4**). The increased number of microglia in the hippocampus of aged mice was prevented in aged *Mlkl-KO* mice, as it was the microglia overlapping with degenerating axons (**Fig S5**), as previously described (Thadathil et al. 2021). Accordingly, measurement of Iba1 mean intensity also indicated that aged *Mlkl*-KO mice present less microglia activation than their WT littermates (**Fig 2d, e**). To confirm the contribution of necroptosis to the inflammatory state of the aging brain, we measured the levels of several cytokines and chemokines in hippocampal lysates and serum of adult versus aged WT and *Mlkl*-KO mice by Luminex High Performance Assay (**Table S1**). Three of the twelve cytokines analyzed showed significant changes under *Mlkl* deficiency in the hippocampus of aged mice. The levels of the pro-inflammatory cytokine IL-12 decreased in aged *Mlkl-KO* hippocampus compared with aged WT mice, reaching levels comparable with adult WT mice (**Fig 2f**). Interestingly, the anti-inflammatory cytokines IL-2 and IL-10 significantly increased in the hippocampus of aged *Mlkl-KO* mice (**Fig 2g**). Moreover, the systemic pro-inflammatory profile also decreased in serum samples from aged *Mlkl-KO* mice (**Fig S6**). These results indicate that necroptosis contributes to brain inflammation by modulating both pro- and antiinflammatory cytokines.

**Fig 2.**
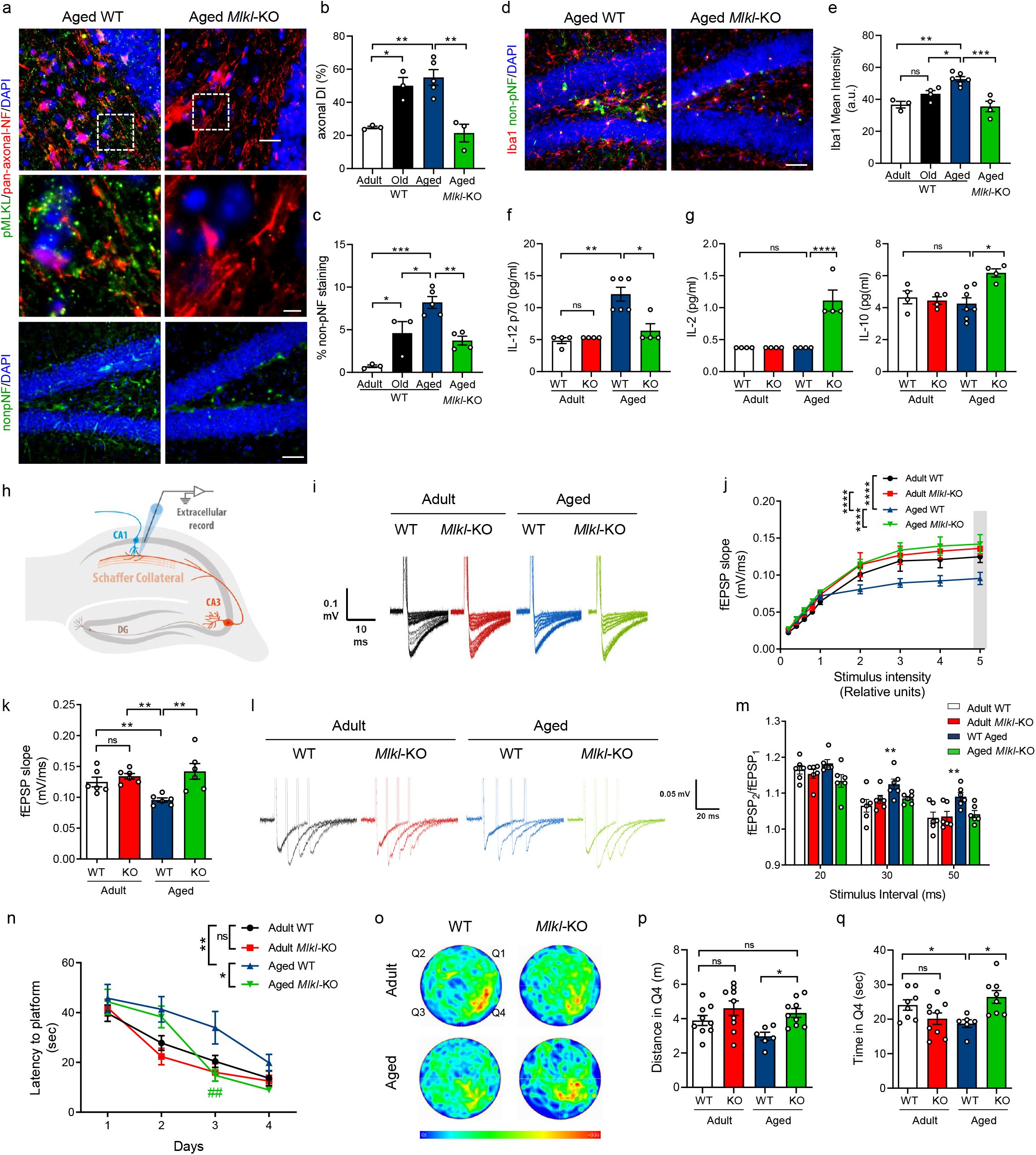
*Mlkl* deficiency protects from neurodegenerative hallmarks and improves hippocampal function and memory in aged mice. Axonal degeneration and neuroinflammation in the hippocampus of aged WT and *Mlkl*-KO mice compared with the normal aging curve (N= 3-5 mice per group). **(a, b)** Axonal degeneration index (DI) calculated from pan-axonal-NF images (red, bars: 25 μm; 5 μm, insets). **(a, c)** Signs of neurodegeneration by non-pNF staining (bar: 50 μm), plotted as percentage of non-pNF area. **(d, e)** Microglia activation quantified as Iba1 mean intensity (bar: 50 μm). **(f, g)** Cytokines levels in the hippocampus of adult vs aged WT and *Mlkl*-KO. Absolute cytokine levels (pg/ml) of IL-12, IL-2 and IL-10 (N=4-7 mice per group). **(h)** Field excitatory post synaptic potential (fEPSP) measured in hippocampal slices in the CA3-CA1 circuit. **(i, j)** Average traces of the evoked potentials plotted as fEPSP slope upon several stimulus intensity. **(k)** Average fEPSP slope at the highest stimulus (gray bar in j). **(l, m)** Paired pulse facilitation as facilitation index (fEPSP_2_/fEPSP_1_), N=3 mice per group (n=3 slices per mice). **(n)** Morris water maze (MWM) navigation task. Escape latency measured as the time to reach the hidden platform during the learning task in aged and adult WT vs *Mlkl-KO* mice. **(o)** Mean heatmaps (pseudo-color) specify location of each mice cohort along time during memory testing (day 5). Q4 quadrant, initial location of the hidden platform during training. **(p, q)** Travelled distance and time spent in Q4. Values, n=7-10 mice per group, mean ± SEM, *p:<0.05; **p<0.01; ***p<0.005; ****p<0.001. One-way ANOVA with Tukey analysis for multiple comparisons. Two-way ANOVA plus Sidak’s multiple comparison test for the learning curve data analysis. *p=0.0209, significance between aged WT and *Mlkl-KO* curves; **p=0.0015, differences between adult and aged WT mice; ^##^p=0.0043, day 3 significance between aged WT vs *Mlkl*-KO.

### 2.3. Loss of MLKL improves hippocampal synaptic transmission, learning and memory in aged mice

To further study whether necroptosis activation affects hippocampal function along aging, we first performed electrophysiological recordings of the CA3-CA1 synapses to evaluate synaptic transmission. Extracellular field-excitatory post-synaptic potentials (fEPSP) were registered in hippocampal slices (**Fig 2h**). An age-dependent decrease in fEPSP slope was observed in WT mice. Nevertheless, in aged *Mlkl*-KO mice, the fEPSP slope was maintained at levels comparable to adult WT mice (**Fig 2i, j, k**). To specifically evaluate whether axonal alterations contribute to an age-dependent decrease in synaptic transmission, we analyzed the facilitation index (fEPSP_2_/fEPSP_1_) using a paired-pulse stimulation protocol (**Fig 2l**). The increased facilitation index observed in WT aged mice indicates a decreased neurotransmitter release probability from the axonal compartment (pre-synapse). Interestingly, the facilitation index of aged *Mlkl-KO* mice was comparable to adult WT mice (**Fig 2m**). These results indicate that *Mlkl* deficiency delays the loss of synaptic strength in the hippocampus inherent to brain aging, mainly preventing axonal function defects in aged mice.

Since memory capabilities depend on proper hippocampal function and both are affected by age (Yang et al. 2019; Burke & Barnes 2006), we evaluated whether *Mlkl* loss improves spatial learning and memory in aged mice using the Morris water maze (MWM) navigation task. The learning curve of aged *Mlkl-KO* mice was significantly faster than those of aged WT animals and reached a reduced latency to find the platform, showing comparable escape time to adult animals (**Fig 2n**). No differences were observed between adult WT and *Mlkl*-KO mice in the learning curve. To evaluate memory, mice were challenged 24 hours after training to find the original location of the platform. Aged *Mlkl*-KO mice travelled a larger distance and spent more time exploring in the target quadrant Q4 compared to aged WT animals (**Fig 2p,q**), without significant changes in the mean swimming speed between both groups (**Fig S7**), demonstrating that loss of *Mlkl* prevents learning and memory loss associated with aging.

Altogether, these results demonstrate that an age-associated increase in brain necroptosis induces degeneration of axons in the hippocampus, thereby depressing synaptic transmission, and impairing hippocampal-dependent functions, such as learning and memory in aged mice.

### 2.4. Pharmacological inhibition of necroptosis reverts key signs of brain aging, improving hippocampal function and memory

To further explore the role of necroptosis in brain aging, we use the RIPK3 inhibitor, GSK’872 (Salvadores & Court 2020; Yang et al. 2017). Diffusion pumps were filled with vehicle or GSK’872, and intraperitoneally implanted to continuously diffuse the inhibitor in 23-month-old mice for 28 days (2 mg/kg GSK’872 at 0.11 μl/hr). Pharmacokinetic studies shown measurable levels of GSK’872 1h post administration in the brain (187.6 ± 17.11 nm) and plasma (18.32 ± 0.85 μm) in aged animals after a single i.p. dose of 10mg/kg. Importantly, GSK’872 treatment leads to decreased pRIPK3 signal in the hilus of aged GSK’872-treated mice (**Fig S8**). Remarkably, aged mice treated with GSK’872 showed decreased non-pNF staining in comparison to vehicle treated mice of the same age (**Fig 3a**), and comparable with those observed in adult animals (**Fig 3b and 2c**). Moreover, decreased microglia activation was also observed, indicating that a short-term treatment with GSK’872 is capable of reverting one of the main signs of brain inflammation associated with aging (**Fig 3c, d**).

**Fig 3.**
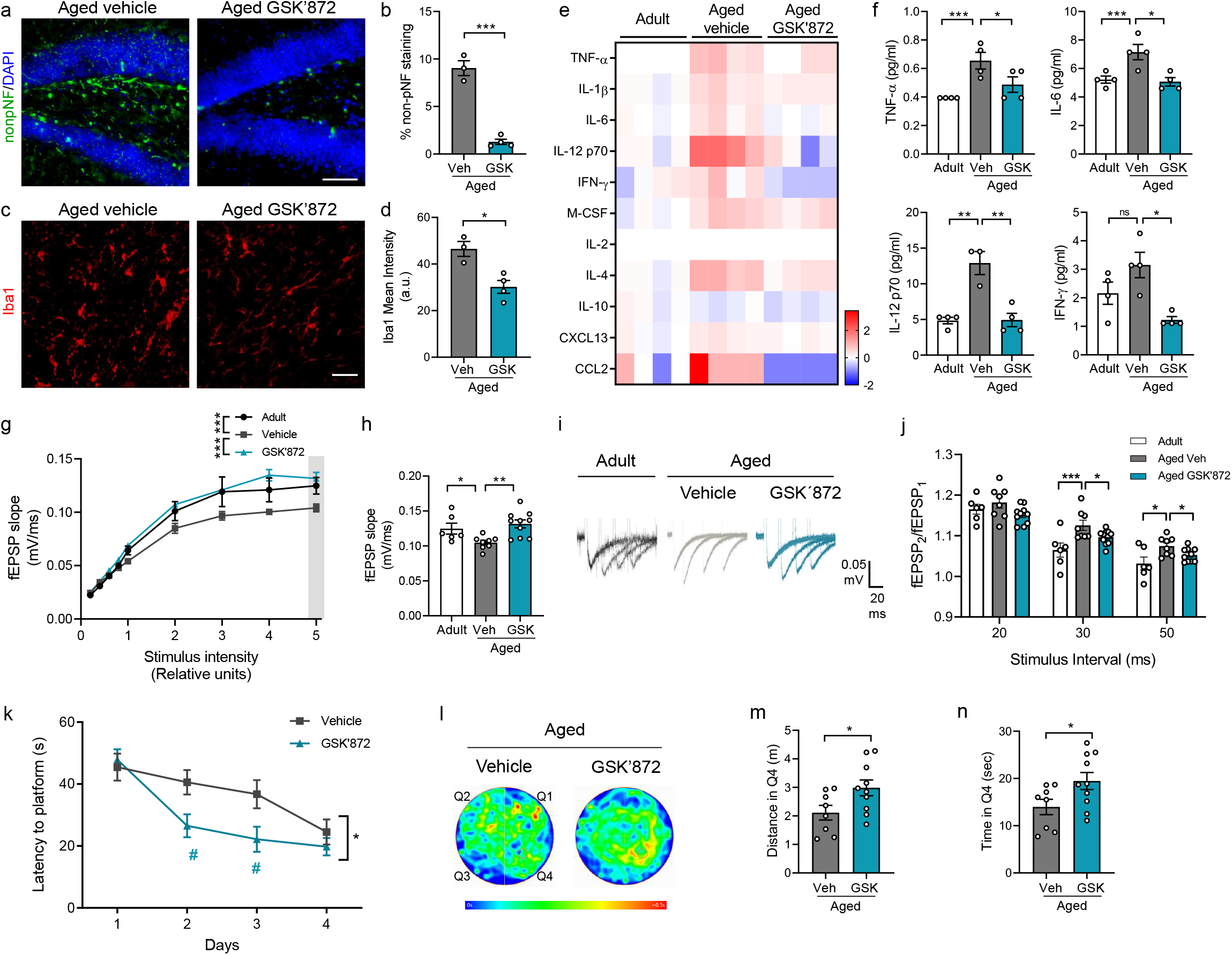
Short-term inhibition of necroptosis reverts hippocampal degeneration restoring electrophysiological function and memory in aged mice. Aged mice treated with vehicle or GSK’872. **(a,b)** Non-pNF staining (bar: 50 μm), plotted as the percentage of stained area. **(c,d)** Microglia activation (bar: 50μm) measured as Iba1 mean intensity. **(e)** Heatmap, fold change levels of cytokines analyzed in the hippocampus (n =4, separated in each column). **(f)** Absolute levels (pg/ml) of TNF-α, IL-6, IL-12p70 and IFN-γ. **(g)** Average traces of the evoked potentials recorded and used to plot the fEPSP slope upon several stimulus intensity. **(h)** Average fEPSP slope from the highest stimulus (gray bar in g). **(i, j)** Facilitation index (fEPSP_2_/fEPSP_1_), N=3 mice per group (n=3 slices per mice). Values, mean ± SEM, *p:<0.05; **p<0.01; ***p<0.005, one-way ANOVA with Tukey analysis for multiple comparisons. **(k)** MWM test, escape latency of vehicle-treated vs GSK’872-treated aged mice. **(l)** Swimming heatmaps (pseudo-color) show average mice location along time during memory testing (day 5). **(m, n)** Travelled distance and the time spent in Q4. Values, n=8-10 mice. T-test with Welch’s correction, *p:<0.01. Two-way ANOVA with Sidak’s multiple comparison test was applied for the learning curve data. *p=0.0259, significance between both curves; #p=0.0210 and p=0.0328, differences between vehicle vs GSK’872 at day 2 and 3, respectively.

Since RIPK3 is involved in pathways that regulate cytokines secretion (Orozco & Oberst 2017) we performed Luminex High Performance Assay to detect changes in a pool of selected cytokines (**Tables S1**). Interestingly, the hippocampus of GSK’872 treated mice showed a pattern of cytokine levels highly similar to those observed in untreated adult mice (**Fig 3e**). The analysis reveals that RIPK3 inhibition mainly reduces pro-inflammatory cytokines, such as TNF-α, IL-6, IL-12, and IFN-γ, reaching youthful-like cytokine levels equivalent to adult mice (**Fig 3f**). A similar profile of decrease in pro-inflammatory cytokines was observed systemically in serum samples of GSK’872 treated mice (**Fig S6**).

At the functional level, electrophysiological recordings in the hippocampus demonstrated equivalent fEPSP between aged mice treated with GSK’872 and untreated adult mice. Average traces showed significant differences between aged vehicle-treated mice versus those that received the RIPK3 inhibitor (**Fig 3g, h**). In addition, facilitation index was significantly lower in the hippocampus of aged mice treated with GSK’872 compared with the aged-vehicle mice (**Fig 3i, j**), showing that late and short-term necroptosis inhibition can revert the loss of synaptic strength in aged mice. Remarkably, RIPK3 inhibition significantly improved learning in aged mice (**Fig 3k**). Memory assessment indicated that aged mice with GSK’872 treatment spent more time in the target quadrant and travelled larger distance in Q4 compared with aged vehicle-treated mice (**Fig 3l-n**). Thus, RIPK3 inhibition is capable to recover aged mice from learning and memory impairment.

These results demonstrate that a short-term systemic administration of GSK’872, in a late phase of the lifespan of mice, can revert key hallmarks of brain aging, including AxD, neuroinflammation, and age-associated memory impairment, proposing the inhibition of necroptosis as an attractive therapeutic target to improve memory in the elderly.

### 2.5. Proteomic analysis in the hippocampus of aged mice with inhibition of necroptosis reveals improvement in key hallmarks of aging and brain rejuvenation

In order to elucidate the specific molecular alterations underpinning our necroptosis-inhibitory approaches toward reducing aging phenotypes, we employed a state-of-the-art single-shot, label free quantitative proteomic approach. Hippocampus of adult and aged WT mice, aged *Mlkl-KO* mice, and aged GSK’872-treated mice were subjected to single-shot label-free mass spectrometry (see workflow in **Fig S9a**), obtaining a high degree of coverage of the proteome with almost 7,000 proteins detected (**Fig S9b**).

After relative expression ratios were calculated and expression profile clustering were performed, we identified subsets of proteins exhibiting opposing directionality in expression between the “normal” aging proteome (aged vs adult animals) and the proteomes of animals subjected to genetic (aged *Mlkl-KO* vs aged WT) or pharmacological (aged GSK’872 vs aged vehicle) inhibition of necroptosis (**Fig S9 and Fig S10**). In doing so, it was possible to reduce the number of correlative candidate proteins to 2,516 proteins whose expression increases and 2,307 which decrease in the aging hippocampus and where genetic and pharmacological modulation leads to a degree of reversion (**Fig 4a**). Pathway analysis confirmed opposing directionality in the activation status of numerous biological processes between “normal” and necroptosis-targeted aged animals. These include several canonical pathways and biological function annotations previously implicated in normal aging (López-Otín et al. 2013; Kennedy et al. 2014) (see **Fig S11a**) and in brain rejuvenation (Bouchard & Villeda 2015; Wyss-Coray 2016) (**Fig 4b and Fig S11b**). Most of the molecular cascades belonging to these pathways are typically associated with neurodegeneration and/or neuronal aging, including molecular cascades involved in *synaptic mechanisms, senescence* and *cellular homeostasis* (**Fig 4b**).

**Fig 4.**
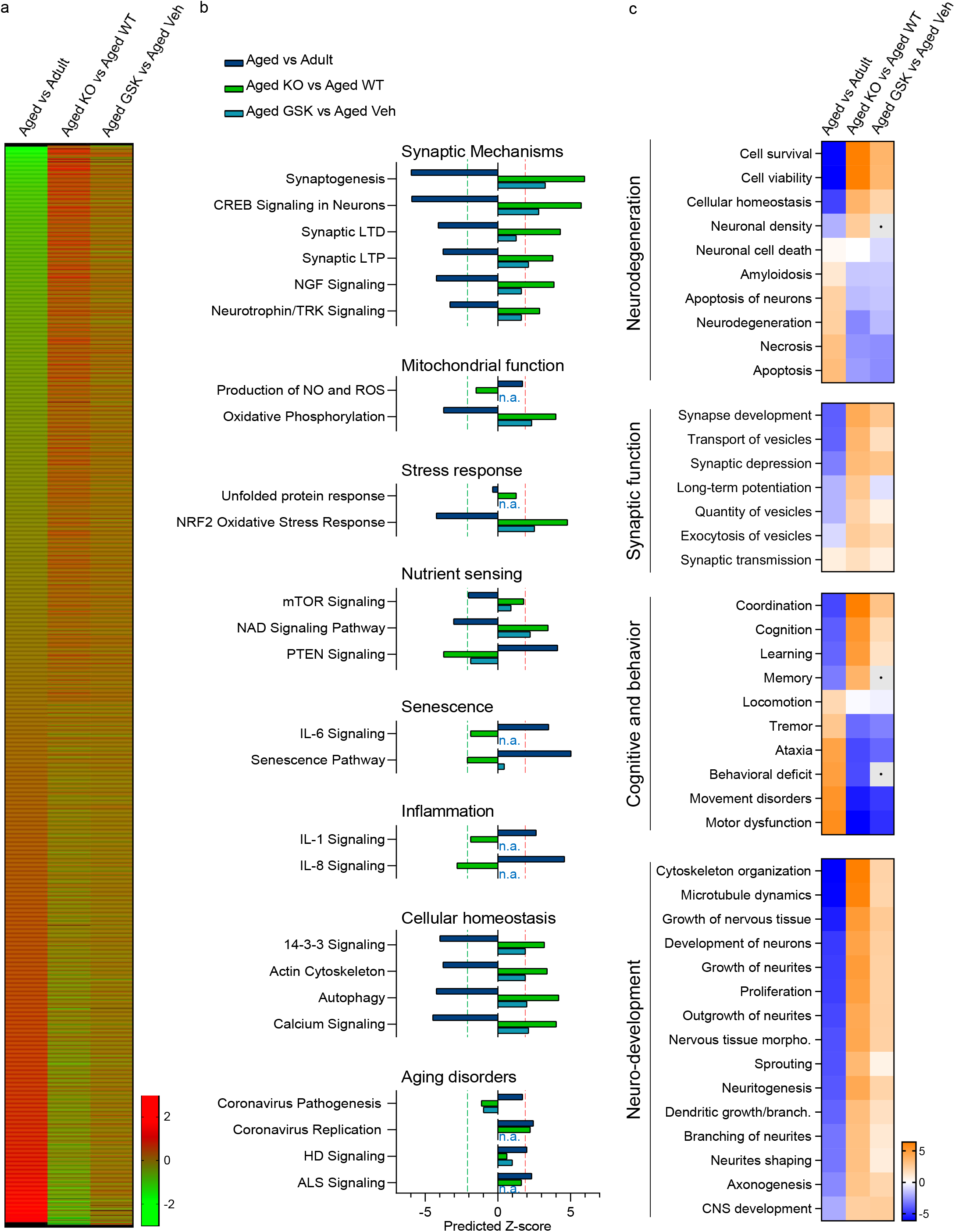
Proteomic analysis reveals opposing directionality in protein expression and aging-associated biofunctions between normal aging and necroptosis-targeted inhibition. **(a)** Heat map, protein expression profile between “normal” aging (aged vs adult) and necroptosis-inhibited processes in aged mice (*Mlkl*-KO and GSK’872). Mean expression ratio (log 2) of *n*=4 mice per experimental group vs their respective controls. Upregulation (red), downregulation (green). **(b)** Canonical pathways classified as aging hallmarks. A predicted z-score >2 or <-2 (indicated by dotted lines) is considered statistically significant. **(c)** Diseases and biofunctions implicated in brain rejuvenation. Heatmaps, predicted activation (orange) and predicted inhibition (blue) scores between “normal” aging, and both *Mlkl-*KO or GSK’872-treated mice.

In order to explore the contribution of necroptosis in age-associated neuronal dysfunction, we evaluated several biological functions affected by normal aging in our proteomic analysis. These biofunctions were classified considering their contribution in central brain functions, designated as *neurodegeneration, synaptic function, cognitive and behavior*, and *neuronal development*. It is interesting to note that the patterns of change of these biofunctions show opposing directionality in both the genetic and pharmacological approaches of necroptosis inhibition compared with normal aging (**Fig 4c**). Remarkably, key molecular and cellular functions associated with neurodegeneration, neuronal integrity and function, showed a clear reversion in the context of pathway analysis in necroptosis-targeted aging in comparison with normal aging (**Fig 4c**). This analysis demonstrated that the inhibition of necroptosis supports proper brain function in aged animals, improving key hallmarks of aging and restoring (in the case of GSK’872) relevant functions involved in brain rejuvenation.

### 2.6. Necroptosis inhibition induces synaptic-long term potentiation in aged mice

Among the cascades elucidated in our proteomic analysis, those classified as *synaptic mechanisms* showed the highest predicted z-score (**Fig 4b**). Synaptic long-term potentiation (LTP) is a key process directly related with learning and memory, and early impaired during aging (Lynch et al. 2006). The contribution of the synaptic LTP signaling at the level of individual molecules is visually illustrated in **Fig 5a.** These molecular changes are prevented in the aging process of the *Mlkl-KO* mice (red molecules) and partially reverted after necroptosis inhibition with GSK’872 in aged animals (**Fig 5b**). We therefore analyzed hippocampal synaptic plasticity by studying LTP magnitude in the CA3-CA1 transmission. By the usage of a high-frequency stimulation protocol, we found that LTP induction was compromised in aged WT mice when compared to adult WT mice (**Fig 5c-e**). Surprisingly, adult *Mlkl*-KO mice presented a higher LTP magnitude than the control WT group and the loss of *Mlkl* in aged mice restored LTP induction and maintenance beyond adult WT mice potentiation. Similar to KO experiments, we detected a reduction in LTP magnitude when we compared adult WT mice with aged vehicle-treated mice. Surprisingly, only one month of GSK’872 treatment in aged mice was capable to improve LTP magnitude, reaching adult-like levels (**Fig 5f-h**). As a correlate of learning and memory improvement, hippocampal long-term synaptic plasticity also impacts dendritic spine remodeling (Engert & Bonhoeffer 1999). We observed that the inhibition of necroptosis in aged *Mlkl*-KO mice protects from the loss of spines in CA1 neurons of the hippocampus along aging (**Fig 5i, j**). Altogether, these results demonstrate that inhibition of necroptosis either by genetic knock-out of *Mlkl* or by pharmacologic RIPK3 inhibition improved or restore synaptic plasticity in aged mice (functionally and morphologically), a synaptic process that is crucial to support brain rejuvenation (Wyss-Coray 2016).

**Fig 5.**
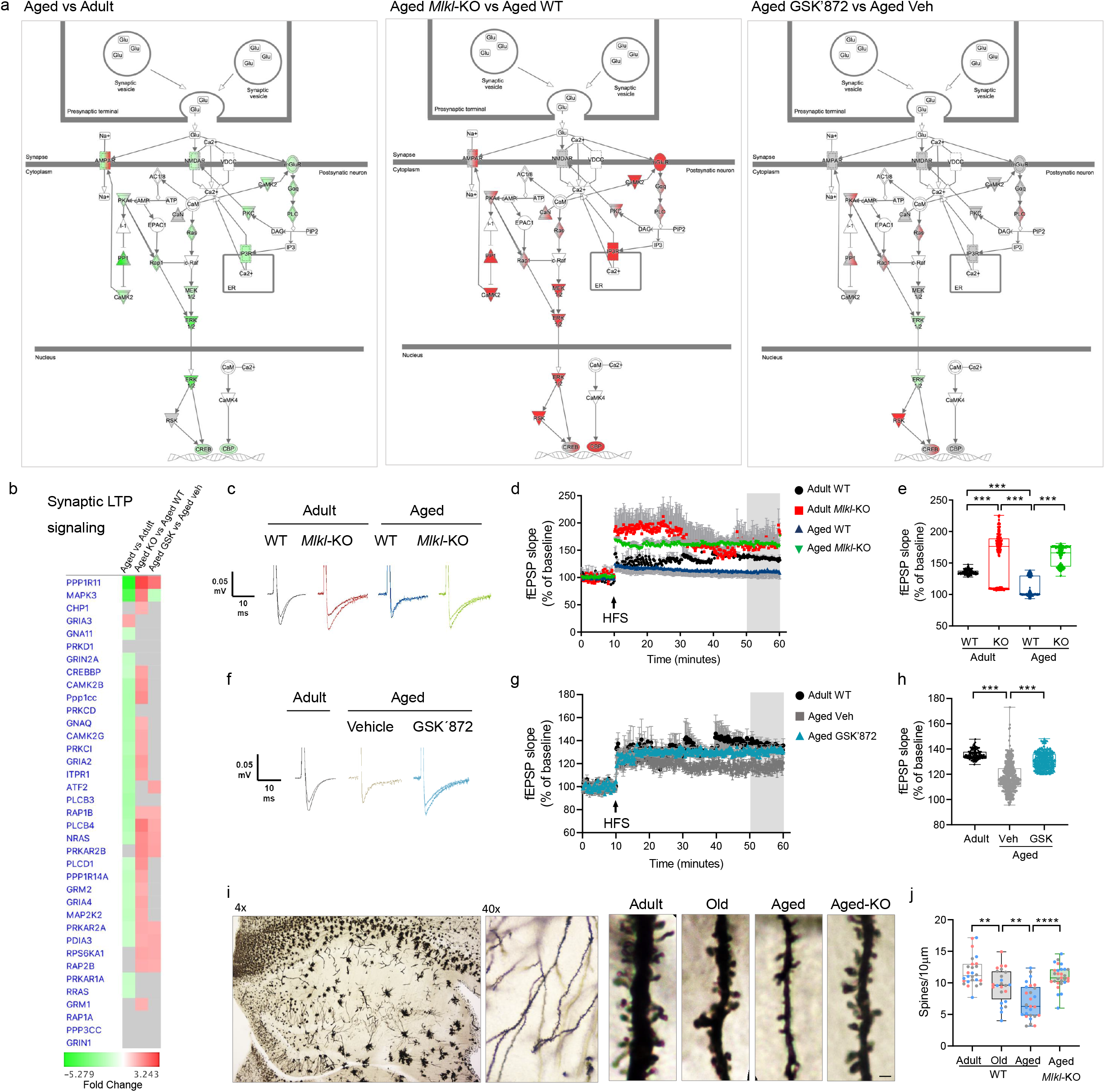
Increased synaptic long-term potentiation in the necroptosis-inhibited aging process. **(a)** Canonical pathway annotation of “Synaptic long-term potentiation signaling”. Upregulation (red) and downregulation (green) compared to control. Molecules in grey fell below the 20% cut-off. Molecules in white were not present within input dataset but changed less than 20% in analysis. Solid connecting lines: direct interaction; dashed connecting lines: indirect interaction. **(b)** Heat map of individual proteins assigned to the “Synaptic long-term potentiation signaling”. Changes were expressed as fold change. **(c-e)** Synaptic plasticity measured as LTP magnitude in the CA1-CA3 hippocampal transmission of WT and *Mlkl-KO* mice from adult and aged groups after high-frequency stimulation (HFS). fEPSP slope plotted as individual values considering the last 10 minutes of recording. **(f-h)** LTP magnitude of aged GSK’872-treated mice vs vehicle, registered for 1h. Boxplots of the last 10 minutes of fEPSP slope. N=4 mice per group (n=8-10 slices per mice). **(i, j)** Dendritic spines visualized by Golgi-Cox staining in the hippocampus along aging and compared with aged *Mlkl-KO* mice (bar: 1 μm). Spines number normalized to 10 μm of dendrite length. (n = 3 mice per group; circles correspond to each dendrite measured per n). Values, mean ± SEM, **p<0.01, ***p<0.005, ****p<0.001. One-way ANOVA.

### 2.7. Necroptosis activation contributes to the acquisition of the age-associated senescent phenotype in the hippocampus during aging

The accumulation of senescent cells in aged tissue, including the brain, is one of the most common feature of aging (Wang et al. 2009; Jurk et al. 2012). As neuronal senescence has also been detected in the hippocampus of aged mice (Gorostieta-Salas et al. 2021), we aimed to determine whether necroptosis contributes to the establishment of the senescent phenotype of hippocampal neurons along aging. From our proteomic data, the molecular changes associated to senescence are mostly prevented in the aging process of the *Mlkl-KO* mice and partially reverted in the pharmacologic inhibition of necroptosis with GSK’872 treatment in aged animals (**Fig 6a**). Interestingly, the expression profile of two key proteins involved in the acquisition of the senescent phenotype, namely CDKN1B and NFκB1 (Pruitt et al. 2013), is mainly reverted in both *Mlkl-KO* and GK’872-treated mice during aging (blue dotted insets, **Fig 6a, b**). We therefore evaluated cell senescence by SA-βgalactosidase (SA-βgal) activity in the hippocampus of aged WT and aged *Mlkl-KO* mice. The accumulation of SA-βgal positive neurons was mainly observed in the CA2/3 subfield of the hippocampus of aged mice (Gorostieta-Salas et al. 2021) (**Fig 6c**), but it was significantly lower in aged *Mlkl*-KO mice (**Fig 6c, d**). A reduced SA-βgal activity was also observed in the hippocampus of aged mice treated with GSK’872 compared with vehicle-treated animals (**Fig 6e, f**), reinforcing the data obtained from the proteomic analysis, which overall indicates that age-related activation of necroptosis contributes to the development of key pathological changes that are involved in brain aging.

**Figure 6.**
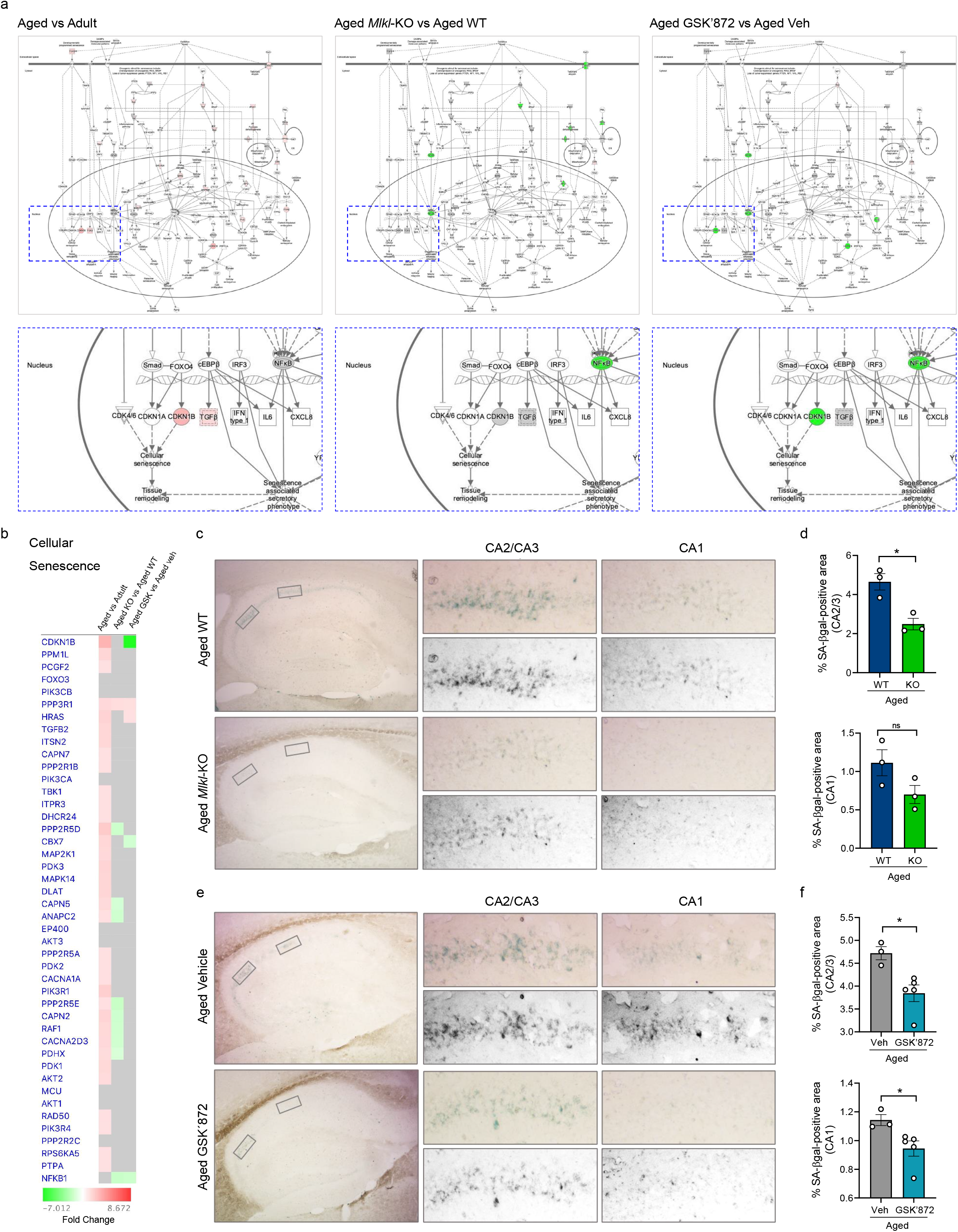
Decreased senescent phenotype in the hippocampus of aged mice with inhibited necroptosis. **(a)** Canonical pathway annotation of “Cellular senescence signaling”. Upregulation (red) and downregulation (green) compared to control. Molecules in grey fell below the 20% cut-off. Molecules in white were not present within input dataset but changed less than 20%. Solid connecting lines: direct interaction; dashed connecting lines: indirect interaction. Magnifications show target molecules involved in cellular senescence execution (blue dotted rectangle) **(b)** Heat map of individual proteins assigned to the “cellular senescence” annotation. Changes were expressed as fold change. **(c, d)** SA-*β*galactosidase (SA-*β*gal) activity quantified in the hippocampus of aged WT vs aged *Mlkl-KO* mice. Magnifications of CA1 and CA2/3 areas show SA-*β*gal staining in grayscale. Percentage of the SA-*β*gal-positive area. **(e, f)** SA-*β*gal activity in the hippocampus of aged vehicle vs GSK’872-treated mice. Values, mean ± SEM (n=3). Unpaired T-test Student, *p<0.05.

## 3. DISCUSSION

Our findings reveal the involvement of necroptosis in the progression of AxD in the hippocampus during normal aging, uncovering relevant implications of necroptosis activation in age-associated cognitive impairment, which might undermine healthy aging. Of note, most detrimental consequences of brain aging, including neuroinflammation, synaptic failure and hippocampal-dependent behavioral impairment were prevented through inhibition of necroptosis by genetic or pharmacological means. In our mouse model of aging, neurodegeneration appears to be restricted to axonal-enriched brain regions, such as the corpus striatum, the cerebellum, and white matter regions of the spinal cord (**Fig S1**). Corresponding with this observation, neurodegeneration was also noted in hippocampal subfields recognized as the richest axonal projection regions, including DG-CA3-CA1 axonal pathways (Ropireddy et al. 2011). The DG supports hippocampal neuronal connectivity by receiving entorhinal cortex information, which is transmitted to the CA3-CA1 neurons to complete the hippocampal circuitry. DG neurons, particularly mossy cells of the hilus, are extremely vulnerable to degenerative insults (Santhakumar et al. 2000; Scharfman 2016), and aging (West 1993), compromising DG connectivity and increasing susceptibility to memory deficits in aged individuals (Amani et al. 2021; Dillon et al. 2017), and aligned with our results showing increased necroptosis in hilar cells, and an early activation in DG hilar fibers along aging (**Fig 1**). These observations suggest that the activation of necroptosis in DG hilar cells and axons may contribute to the vulnerability of this area to degenerate along aging, to adversely impact synaptic function and hippocampal-dependent behavioral performance. The functional role of necroptosis as a crucial pathogenic determinant in experimental models of human diseases has been of increasing interest in the field (Choi et al. 2019). Increased pMLKL levels in aged mice suggests that necroptosis could be also considered as a biomarker of aging progression. Of note, aged *Mlkl*-KO mice presented a youthful phenotype in the hippocampus, reaching levels of AxD comparable with those observed in younger mice (**Fig 2**).

As a chronic inflammatory condition, aging also influences the inflammatory status of the brain (Kennedy et al. 2014), mainly through microglia activation and increase of pro-inflammatory cytokines (Barrientos et al. 2015). Among the pro-inflammatory cytokines analyzed in the hippocampus of aged *Mlkl*-KO mice only IL-12 decreased, reaching levels comparable with adult mice (**Fig 2**). IL-12 is produced in the brain by microglia and required for IFN-γ and TNF-α production, two master inflammatory cytokines. The increased expression of IL-12 in the brain has been associated with spontaneous neurological disorders in aged mice (Hofer et al. 2004). By contrast, the IL12-KO mouse exhibits lower levels of microglia activation and reduced neurodegeneration in an excitotoxicity-mediated injury model (Chen et al. 2004). Additionally, increased levels of the anti-inflammatory cytokines, IL-2 and IL-10, were specifically detected in aged *Mlkl-KO* mice (**Fig 2**), suggesting that MLKL could also act as a repressor of anti-inflammatory cytokine expression under necroptosis activated conditions. Interestingly, IL-10 inhibits the production of IL-12 (Lobo-Silva et al. 2016), which accompanied with the IL-12 decrease observed in the aged *Mlkl-KO* mice generates a positive antiinflammatory feedback loop, limiting neuroinflammation.

It is well documented that neurodegeneration contributes to cognitive decline during normal aging (Bettio et al. 2017). Our electrophysiological analysis demonstrated that loss of *Mlkl* in aged mice produces changes in the hippocampal circuit that prevent the age-dependent loss of synaptic strength (**Fig 2**). The increased facilitation index in aged WT mice is an indicator of a decreased neurotransmitter release probability from the presynaptic compartment. Remarkably, the decreased paired-pulse facilitation index in aged *Mlkl-KO* mice indicates that necroptosis is in fact contributing to these axonal defects in aged mice. Electrophysiological results were supported by improved learning and memory performance in aged *Mlkl-KO* mice, demonstrating that altogether axonal protection, controlled inflammatory status, as well as synaptic transmission restoration in *Mlkl* deficiency favor youthful-like behavior in aged mice. These observations support the notion that lower levels of necroptosis activation improves key cognitive functions that might positively impact the quality of life, favoring healthy aging.

The brain and peripheral organs share common biological mechanisms of aging (López-Otín et al. 2013). Thus, the development of anti-aging drugs that improve cognitive function and hence the quality of life in old age could have a significant potential at improving healthspan. Our pharmacological strategy to inhibit RIPK3 with GSK’872 in aged mice demonstrated to be extraordinarily effective on reverting AxD, neuroinflammation and hippocampus-dependent functional impairment (**Fig 3**). Interestingly, GSK’872 treatment reduced the levels of most of the pro-inflammatory cytokines analyzed in the hippocampus of aged mice, thereby diminishing neuroinflammation. Moreover, the analysis of serum cytokines showed the same regulatory profile as the brain, demonstrating that systemically, the inflammatory condition of aged mice is maintained in a youthful-like state under necroptotic inhibitory terms.

Quantitative proteomic analysis demonstrated that about 7,000 proteins changed their expression profile as a consequence of aging (**Fig 4** and **Fig S9**), of which 2,516 shown to be upregulated and 2,307 were downregulated under necroptosis inhibitory conditions in the hippocampus of aged mice. The pathways analysis unveiled key biological processes with opposed directionality between “normal” and necroptosis-targeted aging, showing that key hallmarks of aging, including synaptic function, mitochondrial dysfunction, stress response, cellular senescence, deficient nutrient sensing, altered metabolism, and others (Kennedy et al. 2014; López-Otín et al. 2013), are positively regulated under necroptosis inhibitory conditions. Interestingly, our pharmacological approach demonstrated a similar trend in effect over the proteome as the genetic model, suggesting that necroptosis inhibition could be an attractive therapeutic strategy to slow aging. In fact, results obtained from our proteomic analysis indicated that GSK’872 treatment influences most of the different hallmarks of aging, an inclusion criteria that is currently recognized to accept novel molecule candidates as geroprotectors (Partridge et al. 2020). Furthermore, inhibition of necroptosis demonstrated ability to modulate several brain functions implicated in brain rejuvenation (Wyss-Coray 2016) (**Fig 4** and **Fig S11**), including those elucidated from the proteomic analysis and experimentally evaluated, such as synaptic plasticity and neuronal senescence (**Fig 5** and **Fig 6**). Moreover, other interesting pathways were also demonstrated to be positively modulated by necroptosis inhibition (**Fig S12-S15**). These pathways include *synaptogenesis signaling, calcium signaling, CREB signaling in neurons*, and others; most of them highly implicated in the maintenance of neuronal homeostasis and functioning, and consequently in brain-dependent functions, including cognitive and behavioral processes (Zia et al. 2021).

Overall, our study demonstrates that necroptosis contributes to the age-associated deterioration of axonal integrity and function, affecting hippocampal synaptic plasticity and neuronal connectivity, and consequently the hippocampal-dependent cognitive function of aged mice. Our results from the pharmacological intervention propose necroptosis inhibition as an interesting and novel therapeutic target to counteract the deleterious effects of aging, thus increasing healthspan and potentially delaying the onset of a range of age-related disabilities.

**MATERIALS AND METHODS (**See Supporting Information for full experimental procedure)

### 3.1. Animals

Wild-type (WT) C57BL/6J male mice of different ages were purchase from the Jackson Laboratory and maintained in the Universidad Mayor animal facility. Aging groups were established as follow: adult (3-6 month), old (12-15 month) and aged mice (more than 20 month). The age range for each mice group was selected in equivalence with the human life phases (Flurkey et al. 2007; Hagan 2017). *Mlkl* knockout mice (*Mlkl*-KO) (C57BL/6 background) were kindly provided by Dr Douglas Green (St. Jude Children’s Research Hospital, Memphis, TN, USA) (Murphy et al. 2013). Genotyping is described as Supporting Information. *Mlkl*-KO mice were reproduced and bred in the Animal Facility of the Sciences Faculty of the Mayor University. Animals were kept under standard conditions of light and temperature and were feed with food and water ad libitum in the Animal Facility of the Sciences Faculty of the Mayor University. The research protocol no. 22–2017 was approved by the Animal Care and Use Scientific Ethic Committee of the Mayor University.

### 3.2. Osmotic pump implantation

Micro-osmotic pumps (Alzet, model 1004) containing the RIPK3 inhibitor GSK’872 (Tocris) (2 mg/kg) were surgically implanted in the peritoneal cavity of 23-month-old mice. The pump allows a constant flux of the drug at 0.11 μl/hr for 28 days. One-month post-surgery, mice (24-month-old) were subjected to behavioral test to evaluate memory and then the tissue was extracted for further analyses.

### 3.3. Immunohistochemistry

Mice were deeply anesthetized with isoflurane and intracardially perfused with isotonic saline followed by 4% paraformaldehyde. Brains were dissected, postfixed overnight in 4% paraformaldehyde at 4 °C, and then incubated in 30% sucrose. Tissue was cryoprotected in optimal cutting temperature compound (OCT, Tissue-Tek) at −20 °C and serial sagittal sections of 20 μm thickness were obtained using a cryostat (Leica, CM1860). Brain sections were pre-mounted on positively charged slides and washed in TBS. After antigen retrieval (80 °C, 30 min, 10 mM citrate buffer, pH 6.0), sections were blocked in TBSB (TBS, 5% BSA and 0.25% Triton X-100), and then incubated overnight at 4 °C with primary antibodies. See Supporting Information for full experimental procedure.

### 3.4. Image Analysis

Fluorescent images were analyzed using the Image J software from the NIH, USA. Axonal degeneration index (DI) was measured in pan-axonal immunostained images as the ratio of the area of fragmented axons against the total axonal area (intact + fragmented particles) in the hippocampus using the particle analyzer algorithm of ImageJ. Fragmented and intact axonal particles were estimated by defining area and circularity of the particles (fragmented: <5 μm^2^ and 0.3 ≤ 1 circularity; intact: ≥ 5 μm^2^ and 0<0.3 circularity) (Arrázola et al. 2019). See **Fig S2** for more details.

Axonal pMLKL analysis was performed in hilar fibers by creating a mask from binarized images of the pan-axonal NF labeling. Threshold was manually defined from the pan-axonal NF satining in the hippocampus of adult mice and then fixed for all the analyses performed. The mean intensity of pMLKL was analyze within the axonal mask area. For non-axonal pMLKL staining, positive cells were counted within a defined area in the hilus and normalized to the total number of cells detected with DAPI.

### 3.5. Learning and memory test

Morris water maze (MWM) navigation task was performed to evaluated spatial memory and learning in adult vs aged WT, *Mlkl-KO* and GSK’872-treated mice. Animals were trained in a 1.2-m-diameter circular pool (opaque water, 50 cm deep) filled with 19-21°C water. A submerged 11.5-cm platform (1 cm below the surface of water, invisible to the animal) was used for training, with a maximum trial duration of 60 s; the mice remained on the platform for 10s at the end of each trial. Each animal was trained to locate the platform for 4 consecutive days following external cues (learning curve), 4 times per day. The test was performed on the fifth day by removing the platform, and swimming was monitored to measure the latency time required to reach the platform and the time spent in each quadrant and in the target quadrant (Q4), and the travelled distance within Q4. Both the learning curve and the test were tracked using an automatic tracking system (ANY-maze video tracking software, Stoelting Co, Wood Dale, IL, USA) to obtain the parameters measured.

### 3.6. Senescence-associated beta-galactosidase (SA-βgal) activity

Histochemical detection of SA-βgal activity were performed as was described before (Debacq-Chainiaux et al. 2009). Briefly, SA-βgal activity was determined by incubation with 1 mg/mL of solution of 5-bromo-4-chloro-3-indolyl β-d-galactopyranoside in 0.04M citric acid/sodium, 0.005M K_3_FeCN_6_, 0.005M K_4_FeCN_6_, 0.15 M NaCl, and 0.002M MgCl_2_ diluted in phosphate-buffered saline (pH 6) for 16h. After the incubation, hippocampal slices were washed with TBS and mounted in superfrost microscope slides (TermoFisher, 6776214) using Fluoromount-G (TermoFisher, 00-4958-02). Images were taken with Nikon Eclipse E200 optic microscope with 4X and 10X objective magnification. ImageJ software was used to process the images. Positive area for SA-βgal activity was measured and representative images are shown.

### 3.7. Statistical Analysis

Statistical significance was stablished at p<0.01 by one-way ANOVA with Tukey’s post-test or two-way ANOVA for multiple comparisons. The analysis was performed using GraphPad Prism Software v8.0 and performed with data obtained from ≥ 3 independent experiments for image analyses and n ≥ 8 animals for behavioral test.

### 3.8. Label-free proteomics

#### S-Trap processing of samples

Samples were processed using S-trap mini protocol (Protifi) (for 310 ug and 110 ug samples) and Strap micro protocol (for low conc samples) as recommended by the manufacturer with little modification. Tryptic peptides were pooled, dried, and quantified using Pierce Quantitative fluorometric Peptide Assay (Thermo Scientific). See Supporting Information for full experimental procedure.

#### LC-MS methods

Samples were injected onto a nanoscale C18 reverse-phase chromatography system (UltiMate 3000 RSLC nano, Thermo Scientific) then electrosprayed into an Q Exactive Plus Mass Spectrometer (Thermo Scientific). The data was acquired using a uPAC-compatible easy spray emitter source operated in positive mode with spray voltage at 2.2 kV, and the ion transfer tube temperature at 275°C. The MS was operated in DIA mode. A scan cycle comprised a full MS scan (m/z range from 345-1155), with RF lens at 60%, AGC target 3E6, orbitrap resolution 70,000, maximum injection time at 200 ms and source fragmentation disabled. The MS survey scan was followed by MS/MS DIA scan events using the following parameters: collision energy mode set to linear with a normalized HCD collision energy set to 25, orbitrap resolution 17500, first fixed mass 200 m/z, AGC target 3E6, maximum injection time 55 ms, isolation windows were variable from 5-66 m/z. The inclusion list (DIA windows) and windows widths are shown in **Table S2**. Data for both MS and MS/MS scans were acquired in profile mode. Mass accuracy was checked before the start of samples analysis.

For data analysis of proteomic data see Supporting Information.

## Supporting information

Graphical Abstract

## Acknowledgments and Funding

This work was supported by grants from the Geroscience Center for Brain Health and Metabolism, FONDAP-15150012 (to FAC and CC), Fondo Nacional de Desarrollo Científico y Tecnológico (FONDECYT) N° 1150766 (to FAC), Michael J Fox Foundation for Parkinson’s Research 17303 (to FAC), Agencia Nacional de Investigación y Desarrollo (ANID) FONDECYT Postdoctorado N° 3180313 (to MSA), BBSRC ISP funding (BBS/E/D/10002071) (to TMW and SLE), BBSRC EastBio (to SI), EMC (to RAK), Centro Científico y Tecnológico Ciencia & Vida, FB210008, Financiamiento Basal para Centros Científicos y Tecnológicos de Excelencia de ANID (to SB), Fondo Nacional de Desarrollo Científico y Tecnológico (FONDECYT) N° 1190620 (to WC), Center for Excellence in Science and Technology AFB 170005, PFB 12/2007 (to WC), Fondo Nacional de Desarrollo Científico y Tecnológico (FONDECYT) N° 1200255 to CC, and a PhD fellowship by ANID (to ML).

## Conflict of Interest statement

Authors have no conflicts of interest to disclose.

## Data availability statement

Raw data from Fig 1,2,3,4,5 and 6 are available as an extended excel file named “Figures Raw Data”.Raw data from Supplementary Figures S1, S3, S4, S5, S6, S7 and S8 are available as an extended excel file named “Suppl Figures Raw Data”. The datasets generated from the proteomic analysis are available in the data repository of Edinburgh University with an assigned DOI. All the data that support the findings of this study are also available from the corresponding author upon request.

## Author contributions

MSA, RAK, GU, SB, JCC, WC, TMW and FAC designed experiments, analyzed data, and wrote the manuscript; MSA, ML, GQ, FV, DJL and HH performed experiments; SI, SLE, RAK and TMW performed bioinformatic and proteomic analysis.

## Notes

### Competing Interest Statement

The authors have declared no competing interest.

### Summary of Updates

Author affiliations updated title changes new abstract manuscript modifications figures were regrouped: Fig1: former fig1+fig2 Fig2: former fig3+fig4 Fig3: former fig5+fig6 Fig4: former fig7 Fig5: former fig8 + new data Fig6: Former fig 9 A graphical abstract was added

