## Supplementary material for "Necroptosis inhibition counteracts neurodegeneration, memory decline and key hallmarks of aging, promoting brain rejuvenation": Supplemetary Information

*Running title: Necroptosis inhibition prevents brain aging (*40 characters)

Macarena S. Arrázola[1,2], Matías Lira[3], Gabriel Quiroz[2], Felipe Véliz-Valverde[1,2], Somya Iqbal[4], Samantha L Eaton[4], Rachel A Kline[4], Douglas J Lamont[4], Hernán Huerta[1,2], Gonzalo Ureta[5], Sebastián Bernales[5], J César Cárdenas[1,2,6,7], Waldo Cerpa[3,8], Thomas M. Wishart[4] and Felipe A. Court [1,2,6,*].

[1] Center for Integrative Biology, Faculty of Sciences, Universidad Mayor

[2] Geroscience Center for Brain Health and Metabolism (GERO), Santiago, Chile.

[3] Departamento de Biología Celular y Molecular, Facultad de Ciencias Biológicas, Pontificia Universidad Católica de Chile, Av. Libertador Bernardo O´Higgins 340, Santiago, Chile.

[4] The Roslin Institute, University of Edinburgh, Edinburgh, UK.

[5] Fundación Ciencia & Vida, Santiago, Chile.

[6] Buck Institute for Research on Aging, Novato, CA, USA.

[7] Department of Chemistry and Biochemistry, University of California, Santa Barbara, California, USA

[8] Centro de Excelencia en Biomedicina de Magallanes (CEBIMA), Universidad de Magallanes, Punta Arenas, Chile.

* Contact information:

### 1. SUPPLEMENTARY MATERIAL AND METHODS

#### 1.1. *Mlkl*-KO mice genotyping

Mice were genotyped by standard polymerase chain reaction methods using a primer mix (5'-TAT GAC CAT GGC AAC TCA CG-3', 5'-ACC ATC TCC CCA AAC TGT GA-3' and 5'-TCC TTC CAG CAC CTC GTA AT-3') that distinguished between wild-type (498 bp product) and the recombined Δ-exon 3 (158 bp) Mlkl loci (Murphy et al. 2013).

#### 1.2. Quantification of GSK'872 in brain and plasma

Brain and plasma samples were obtained from aged mice treated for 1h with GSK'872 (10mg/kg i.p.). The bioanalysis of plasma and brain samples was conducted by LC-MS/MS with a QTRAP 4500 triple quadrupole mass spectrometer (Applied Biosystems SCIEX) in the negative ion mode and interfaced with an Ekspert ultraLC 100-XL UHPLC System (Eksigent). Calibration standards (0.003 to 10 μM) and quality controls (0.02, 0.2 and 2.0 μM) were prepared from naïve mouse plasma in parallel with mouse plasma study samples (60 μL) by precipitation with three volumes of ice-cold acetonitrile containing 20 μM of theophylline. The precipitated samples were centrifuged at 6,100 g for 30 min at 4°C. Following centrifugation, an aliquot of each supernatant was transferred to an autosampler vial and diluted with two volumes of aqueous mobile phase (0.2% formic acid in water). Samples were injected onto a reverse phase analytical column (YMC Triart C18; 2.0 x 50 mm; 1.9 μm; YMC CO) and eluted with a gradient of 0.2% formic acid in Acetonitrile. GSK'872 was monitored by a multiple reaction monitoring (MRM) experiment using an Analyst software (v1.6.2, Applied Biosystems SCIEX). Quantitation was conducted using a MultiQuant software (v2.1, Applied Biosystems SCIEX) and the resulting calibration curve was fitted with a linear regression and 1/x weighting. The lower limit of quantitation (LLOQ) was 0.010 μM.

#### 1.3 Immunohistochemistry

Primary antibodies: pMLKL (1:200, phospho S345, Abcam, ab196436), neurofilament pan-axonal cocktail (1:250, SMI312 clone, Biolegend), non-phosphorylated neurofilament (1:250, SMI32 clone, Biolegend), and Iba1 (1:500, Wako Chemicals, 016-26721). Sections were washed to remove the excess of primary antibody and incubated with the appropriate Alexa-Fluor secondary antibodies for 2 h at 25 °C (1:1000, Thermo Fisher Scientific). The excess of secondary antibody was washed, and the sections were coverslipped with Fluoromount-G antifade mounting medium (EMS, 17984-25) containing DAPI staining for nuclei detection (Thermo Fisher Scientific). The anti-pMLKL antibody purchase from Abcam (ab196436) was KO-validated by comparison of the immunodetected signal in aged WT mice with the *Mlkl*-KO mice in the hippocampus (**Fig S16**). Immunostained sections were scanned in a Leica DMi8 Fluorescence Microscope fully automatized.

#### 1.4 Histological Analysis

Free-floating sections were processed for immunohistochemistry as previously described(Oñate et al. 2020). Briefly, slices were quenched with 0.3% $H_2O_2$ for 30 min, blocked with 5% BSA and 0.2% triton X-100 for 2 h and incubated with primary antibody (rabbit anti-pMLKL, 1:200 Abcam) overnight at 4 °C. Then, sections were washed with 0.1 M PBS and incubated with secondary biotinylated antibody (goat anti-rabbit, 1:500 Vector Laboratories) for 2 h at RT. After washing, slices were incubated with avidin–biotin–peroxidase complex (Vector Laboratories) for 1 h at RT followed by 0.1 M PBS washes and developed with 3,3-diaminobenzidine (DAB, Sigma-Aldrich). Finally, sections were co-stained with Nissl (cresyl-violet staining) to detect nuclei and with Eriochrome-C staining for myelinated-axons detection, and finally mounted on glass slides with Entellan medium (Merck).

##### 1.5 Fluoro-Jade C staining

Brain tissue was mounted on positively charged slides and rehydrated on decreasing concentrations of ethanol. After rehydration, the tissue was pre- treated for 10 min in potassium permanganate and then incubated for 10 min in the dark at 25 °C in Fluoro-Jade C and DAPI (Biosensis, TR-100-FJ). The tissue was then washed with water and let dry overnight. The next day the slides were cleared in xylene and coverslipped with DPX-new mounting solution (Merck Millipore).

##### 1.6 Electrophysiology

Electrophysiological recordings were performed as described before (Carvajal et al. 2018). Briefly, transverse slices (400 μm) from the dorsal hippocampus were cut under cold artificial cerebrospinal fluid (ACSF, in mM: 124 NaCl, 2.6 $NaHCO_3$, 10 D-glucose, 2.69 KCl, 1.25 $KH_2PO_4$, 2.5 $CaCl_2$, 1.3 $MgSO_4$, and 2.60 $NaHPO_4$) using a Vibratome (BSK microslicer DTK-1500E, Ted Pella, Redding, CA, USA) and incubated in ACSF for 1 hour at room temperature. In all experiments, 10 μM PTX was added to suppress inhibitory GABAA transmission. Slices were transferred to an experimental chamber (2 ml), superfused (3 ml/min, at room temperature) with gassed ACSF (using 95% $O_2$/5% $CO_2$) and visualized by trans-illumination with a binocular microscope (Amscope, Irvine, CA, USA). To evoke field excitatory post synaptic potentials (fEPSPs), Schaffer collaterals were stimulated with bipolar concentric electrodes (Tungsten, 125 μm OD diameter, Microprobes) connected to an isolation unit (Isoflex, AMPI, Jerusalem, Israel). The stimulation was performed in the *stratum radiatum* within 100–200 μm from the recording site. Recordings were filtered at 2.0–3.0 kHz, sampled at 4.0 kHz using an A/D converter (National Instrument, Austin, TX, USA), and stored with the WinLTP program. The basal excitatory synaptic transmission was measured using an input/output curve protocol with 10 s of interval between stimuli. Data were collected and analyzed offline with pClamp 10 software (Molecular Devices, San Jose, CA, USA). To generate LTP, we used high-frequency stimulation (HFS) protocol, which consisted of 3 trains at 100 Hz of stimuli with an inter-train interval of 10 s. Data were collected and analyzed offline with pClamp 10 software (Molecular Devices, San Jose, CA, USA).

##### 1.7 Golgi-Cox staining

Golgi-Cox impregnation method was used to analyze dendritic spine density in hippocampal slices by using the FD Rapid GolgiStain™ Kit, following manufacturer instructions (FD Neurotechnologies Inc, MD, USA). See detailed protocol in Supporting Indormation. Briefly, fixed brains by PFA perfusion were immersed in impregnation solution (Solution A/ B), and store at room temperature for 2 weeks in the dark. After 72 h in precipitation Solution C, brains were quickly frozen with dry ice and immediately sectioned with a cryostat at a thickness of 120 μm. Coronal brain sections were mounted on gelatin-coated slides, stained with solution D/E, dehydrated with sequential rinses of 50%, 75%, 95% and 100% ethanol and finally mounted with Entellan medium (Merck). Dendritic spines were imaged as Z-stacks images with a Nikon Eclipse E200 microscope by using a 100X objective with immersion oil. Image analyses were performed with the Image J software by Z-projection of the stacks (sum stacks) followed by the measurement of dendrite length with the segmented line tool. Dendritic spines were manually counted in the defined dendrite length and plotted as spine number normalized to 10 µm of dendrite. Between 8 and 10 dendrites were imaged and quantified per mouse, considering n=3 mice per experimental group.

**1.8 Luminex Assay**

Cytokines levels were analyzed by Luminex Mouse Discovery Assay (R&D Systems, MN, USA) using a self-designed panel of 12 selected cytokines (plate code: LXSAMSM-12), based on color-coded beads, pre-coated with analyte-specific capture antibodies that permits simultaneous analysis of the analytes. Table S1 detailed cytokines and chemokines of the panel, bead region, sensitivity, and the main functions. Analysis and detection were performed in a Dual-laser flow-based detection instrument, Luminex 200 analyzer by Proyecto Luminex, Programa de Virología, Redeca, ICBM, Facultad de Medicina, Universidad de Chile.

1.9 **Label-free proteomics**

*S-Trap processing of samples*

Samples were processed using S-trap mini protocol (Protifi) (for 310 ug and 110 ug samples) and S-trap micro protocol (for low conc samples) as recommended by the manufacturer with little modification. After, application of the samples on the S-trap mini spin column, trapped proteins were washed 5 times with S-TRAP binding buffer. A double digestion with trypsin (1:40) was carried out first overnight at 37$^{o}$C in TEAB at a final concentration of 50 mM, and then for another 4 hrs (1:40) in 50mM TEAB. Elution of peptides from S-trap mini spin column was achieved by centrifugation at 1000 x g for 1 min by adding 50 mM TEAB, then 0.2% aqueous formic acid and finally 50% acetonitrile/0.2% formic acid. Resulting tryptic peptides were pooled, dried, and quantified using Pierce Quantitative fluorometric Peptide Assay (Thermo Scientific).

*LC-MS methods*

1.5 µg peptide was analysed per sample. Samples were injected onto a nanoscale C18 reverse-phase chromatography system (UltiMate 3000 RSLC nano, Thermo Scientific) then electrosprayed into an Q Exactive Plus Mass Spectrometer (Thermo Scientific). For liquid chromatography buffers were as follows: buffer A (0.1% formic acid in Milli-Q water (v/v)) and buffer B (80% acetonitrile and 0.1% formic acid in Milli-Q water (v/v). Sample were loaded at 10 µL/min onto a trap column (100 µm × 2 cm, PepMap nanoViper C18 column, 5 µm, 100 Å, Thermo Scientific) equilibrated in 0.1% trifluoroacetic acid (TFA). The trap column was washed for 5 min at the same flow rate with 0.1% TFA then switched in-line with a µPAC C18 nano-LC column (200 cm, inter-pillar distance- 2.5 µm, pore size- 100-200 Å, PharmaFluidics). The peptides were eluted from the column at a constant flow rate of 300 nl/min with a linear gradient from 3.8% buffer B to 12.5% buffer B in 22 mins, then from 12.5% buffer B to 41.3% buffer B in 95 mins, then from 41.3% buffer B to 61.5% in 23 mins and finally to 100% buffer B in 10 mins. The column was then washed with 100% buffer B for 10 min and re-equilibrated in 1% buffer B for 38 mins. Two blanks were run between each sample to reduce carry-over. The column was kept at a constant temperature of 50$^{o}$C. The data was acquired using a uPAC-compatible easy spray emitter source operated in positive mode with spray voltage at 2.2 kV, and the ion transfer tube temperature at 275$^{o}$C. The MS was operated in DIA mode. A scan cycle comprised a full MS scan (m/z range from 345-1155), with RF lens at 60%, AGC target 3E6, orbitrap resolution 70,000, maximum injection time at 200 ms and source fragmentation disabled. The MS survey scan was followed by MS/MS DIA scan events using the following parameters: collision energy mode set to linear with a normalized HCD collision energy set to 25, orbitrap resolution 17500, first fixed mass 200 m/z, AGC target 3E6, maximum injection time 55 ms, isolation windows were variable from 5-66 m/z. The inclusion list (DIA windows) and windows widths are shown in **Table S2**. Data for both MS and MS/MS scans were acquired in profile mode. Mass accuracy was checked before the start of samples analysis.

##### 1.10. Analysis of proteomic data

*Data filtering and generation of expression ratios*

Raw data files from single-shot label-free experiments were converted into Microsoft Excel workbooks and utilised to generate ratios of protein expression within each animal relative to mean expression of n=4 control animals within each comparison (ie. adult: adult (expression ratio=1), aged: adult, *Mlkl*-KO: aged-WT; aged GSK'872: aged vehicle). Proteins identified by fewer than 2 unique peptides were excluded from subsequent analyses in order to ensure maximum identification confidence (**Figure S9**). Relative expression ratios per study (ie. aged vs. adult, *Mlkl*-KO vs. aged WT, and aged GSK'872 vs. aged vehicle) were used for subsequent expression profile clustering analyses. UniProt Accession numbers of proteins identified by 2 or more unique peptides with accompanying expression ratios generated as described above were subjected to expression profile clustering in *BioLayout Express$^{3D}$*. *BioLayout* utilises a user-determined Pearson correlation and the Markov Clustering Algorithm to cluster input data based on user-determined parameter(s) (Enright 2002; Theocharidis et al. 2009). Pearson correlation was set to 0.97 to cluster datasets into distinct subsets based on similarity in expression profile. Discrete clusters exhibiting biologically relevant expression profiles- ie. opposing directionality between aged versus both MLKL-KO and GSK'872 expression ratios (**Fig S10**) were identified and exported as .txt files containing and identifier column and expression ratios, for subsequent analyses in IPA. See Supporting Information for detailed procedure.

*Ingenuity Pathway Analysis (IPA)*

The Ingenuity Pathway Analysis (IPA) application (Ingenuity Systems, Silicon Valley, CA) was used to visualise and explore the cellular and molecular pathways that may have been altered as result of genetic (*Mlkl*-KO) or pharmacological (GSK'872) inhibition of necroptosis. Without user-directed manipulation, IPA's statistical predictions and annotations are approximately 90% based off on peer-reviewed publications; the remaining 10% of stored interactions have been identified by other in silico techniques. The analyses were performed only using experimentally reported interactions published in peer-reviewed publications stored within the "hand-curated" and continually updated Ingenuity Knowledge database (Ingenuity Systems, Silicon Valley, CA). For more information on the computational methodology underpinning IPA, please refer to http://www.ingenuity.com/.

Prior to all analyses within IPA, input datasets comprising, as described above, mean expression ratios of *n*=4 animals per experimental group, were converted to fold-change values, and a ±20% cut-off in expression change respective to control was applied within each respective study. Individual analyses of aged vs. adult, *Mlkl*-KO vs. WT, and GSK'872 vs. vehicle were performed prior to a comparative analysis in order to gain insight into potential biological networks distinguishing "normal" versus "necroptosis-inhibited" aging processes.

For canonical pathway analysis, p-values of canonical pathway scores and subsequent ranking for all analyses performed in this study were derived from a Fisher's Exact Test calculating overlap between molecules in each respective input dataset and number of molecules comprising canonical pathway as defined by the Ingenuity Systems Database. Predicted activation z-scores were calculated by weighing the predicted expression change of target molecules as defined by Ingenuity Knowledge Database against the actual expression change of target molecules reported in input dataset. An activation z-score >2 or <−2 is considered statistically significant (Ingenuity Systems, Silicon Valley, CA). Constituent molecules within pathway were colourized with intensity of colour corresponding to magnitude of change.

For diseases and functions analysis, predicted activation z-scores of associated downstream diseases and functions were calculated by weighing the predicted expression change of target molecules associated with specific "diseases or functions" annotation as defined by Ingenuity Knowledge Database against the actual expression change of target molecules reported in input datasets. An activation z-score >2 or <−2 is considered statistically significant. *P*-values of overlap is derived from a Fisher's exact test were derived from a Fisher's Exact Test calculating overlap between molecules in each respective input dataset and number of molecules comprising the known interactome of each regulator as defined by Ingenuity Systems Database. In graphical format, target molecules present within each proteomic dataset predicted to be activated or inhibited to mediate the associated "diseases or functions" annotation were visualised in relation to their associated predicted regulator and were colourised with intensity of colour corresponding to magnitude of change.

#### 2 SUPPLEMENTARY FIGURES

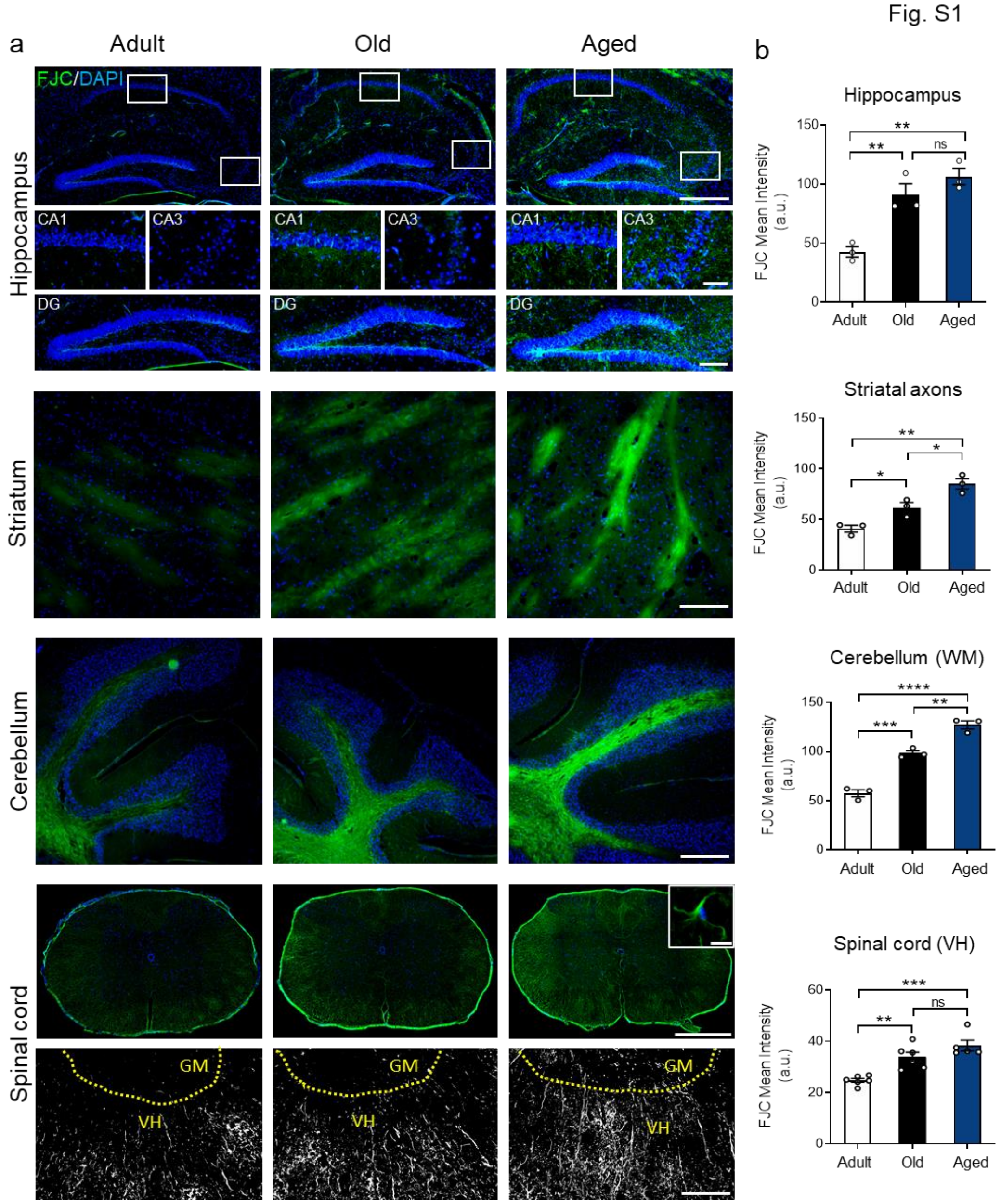


**Fig S1. Progression of neurodegeneration in different brain regions along aging.** Representative images of Fluoro Jade C (FJC, green) staining in the hippocampus (calibration bar, 500 µm; CA1/3 and DG bars are 100 µm and 200 µm respectively), striatum (calibration bar, 25 µm), cerebellum (calibration bar, 50µm) and spinal cord (calibration bar, 100 µm and 25 µm for the inset) of adult (3-6 month), old (12-15 month) and aged mice (more than 24 month). DAPI staining (blue) detects nuclei **(a)**. Mean intensity of FJC was evaluated in the entire hippocampus, the axonal tracts of the striatum, the cerebellar white matter, and in the ventral (VH) of the spinal cord, GM: gray matter **(b)**. Values are the result of the analysis of n=3-4 mice per group. One-way ANOVA with Tukey analysis for multiple comparisons, *p:<0.05; **p<0.01; ***p<0.005; ****p<0.001.

Fig. S2

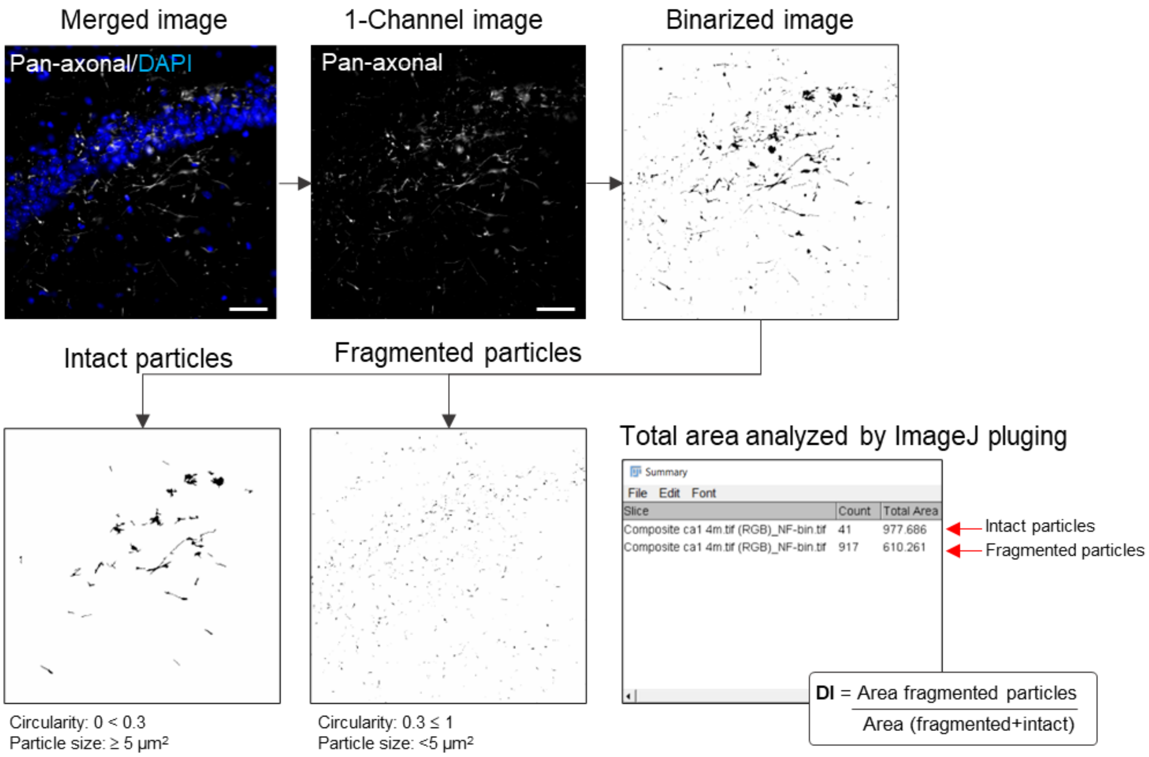


**Fig S2. Workflow of Axonal Degeneration Index analysis.** Raw images of pan-axonal NF labeling were binarized and subjected to the “Analyze Particles” tool of ImageJ software. Fragmented and intact axonal particles were estimated by defining area and circularity of the particles (fragmented: <5 μm$^2$ and 0.3 ≤ 1 circularity; intact: ≥ 5 μm$^2$ and 0<0.3 circularity). Calibration bar, 10 μm. Degenereation Index (DI) was calculated as the ratio betweeem the area of fragmented axons over the total axonal area.

Fig. S3

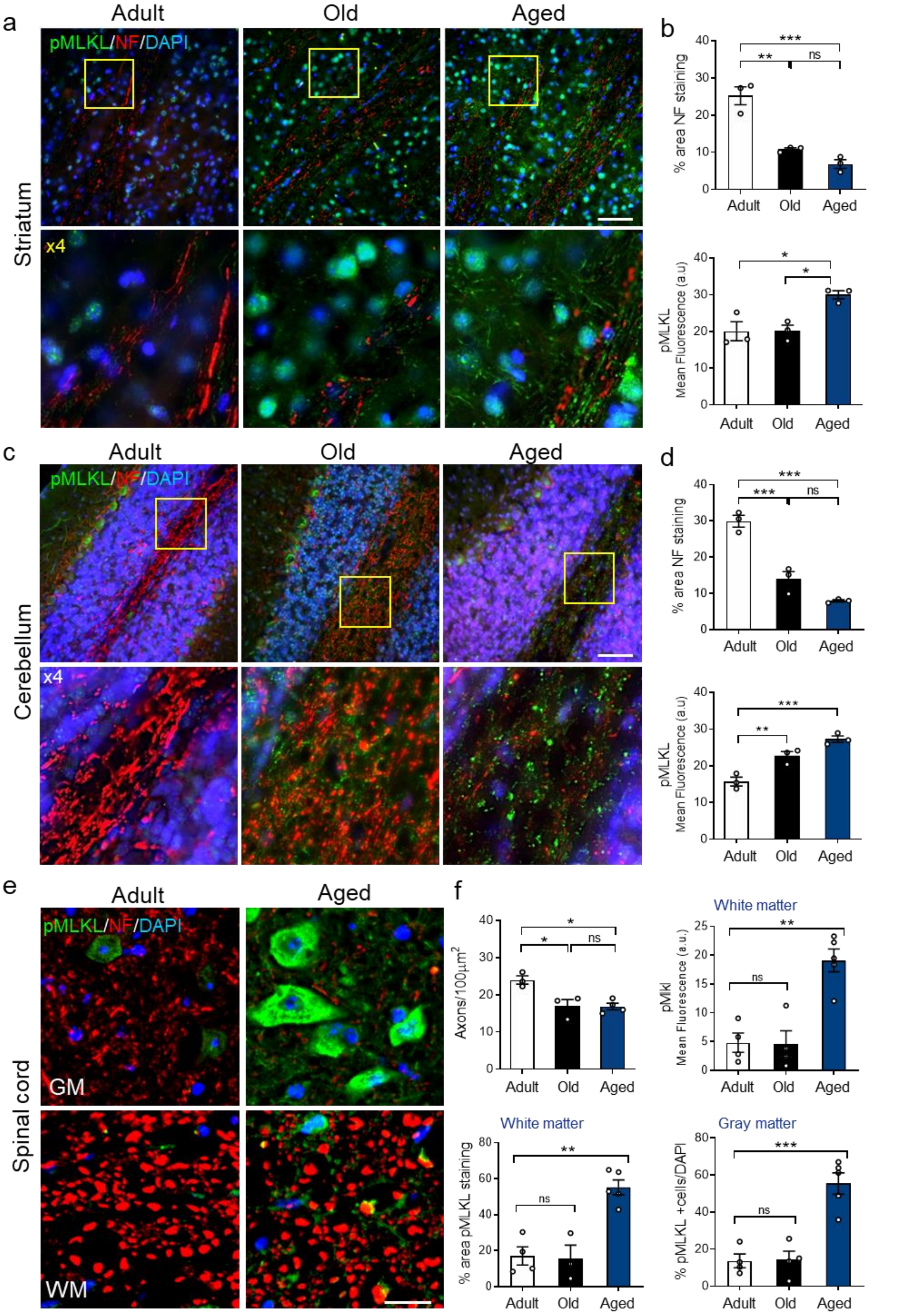
a
Adult
Old
Aged
pMLKL/NF/DAPI
Striatum
x4
b
% area NF staining
***
**
ns
Adult
Old
Aged
pMLKL
Mean Fluorescence (a.u)
*
*
Adult
Old
Aged
c
Adult
Old
Aged
pMLKL/NF/DAPI
Cerebellum
x4
d
% area NF staining
***
***
ns
Adult
Old
Aged
pMLKL
Mean Fluorescence (a.u)
***
**
Adult
Old
Aged
e
Adult
Aged
pMLKL/NF/DAPI
Spinal cord
GM
WM
f
Axons/100μm2
*
*
ns
Adult
Old
Aged
White matter
pMlkl
Mean Fluorescence (a.u.)
**
ns
Adult
Old
Aged
White matter
% area pMLKL staining
**
ns
Adult
Old
Aged
Gray matter
% pMLKL +cells/DAPI
***
ns
Adult
Old
Aged

**Fig S3. Necroptosis activation is correlated with axonal degeneration in diverse areas of the aging brain.** Necroptosis activation was measured through the phosphorylation of MLKL (pMLKL in green) in axon-enriched subfields of different brain areas. Pan-axonal NF antibody (red) was used to stain axons, and DAPI (blue) to detect nuclei and to identify granular cell layers and gray matter regions in the striatum, cerebellum and spinal cord, respectively. **(a,b)** Phosphorylated-MLKL mean intensity increases in axonal tracts of the striatum along aging. Decreased pan-axonal NF staining was used as a readout of axonal degeneration (calibration bar, 25 µm). **(c,d)** Pan-axonal NF staining decreased in the white matter (WM) of the cerebellum, accompanied by an increase in pMLKL staining in the same compartment along aging (calibration bar, 25 µm). **(e,f)** Phosphorylated MLKL levels were evaluated in the white matter (WM) and gray matter (GM) of spinal cord cross-sections during aging. Necroptosis was slightly detected in motor neurons (GM) of adult mice, while no pMLKL signal was observed in the WM (calibration bar, 20 µm). Aged mice present high levels of pMLKL in both, WM and GM measured as pMLKL mean intensity and percentage of stained area. Percentage of pMLKL-positive cells (normalized to DAPI) in the GM also increased in aged mice compare to adult and old groups. Axonal degeneration was assessed by counting the number of axons per area in the WM. Values are the result of the analysis of n=3-6 mice per group. One-way ANOVA with Tukey analysis for multiple comparisons, *p:<0.05; **p<0.01, ***p<0.005.

Fig. S4

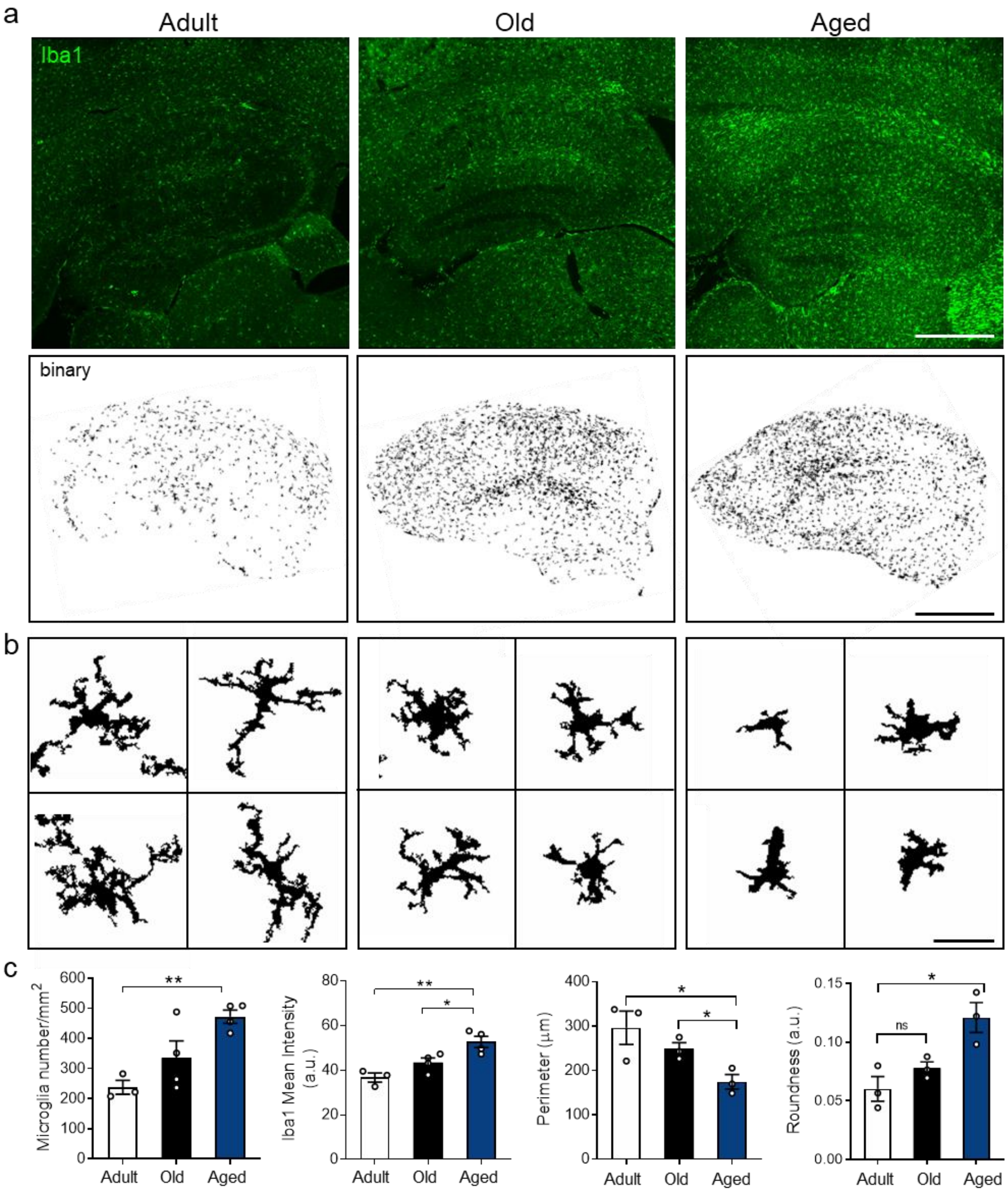


**Fig S4. Increased neuroinflammation is correlated with morphological changes associated to pathological microglia activation in aging. (a)** Microglia activation and cell number were evaluated by Iba1 immunostaining (green), (calibration bar, 200 µm). **(b)** Morphological classification of microglia was performed from Iba1-stained and binarized images. (Adult: resting microglia; Old: primed microglia; Aged: pathologic activation, calibration bar, 30 µm). **(c)** Increased number of microglia and Iba1 intensity were observed in the hippocampus of aged mice. Morphological parameters of microglia reshaping during activation were also evaluated, including decreased perimeter and cell roundness. Values are the result of the analysis of n=3-8 mice per group. One-way ANOVA with Tukey analysis for multiple comparisons, *p:<0.05. **p<0.01.

Fig. S5

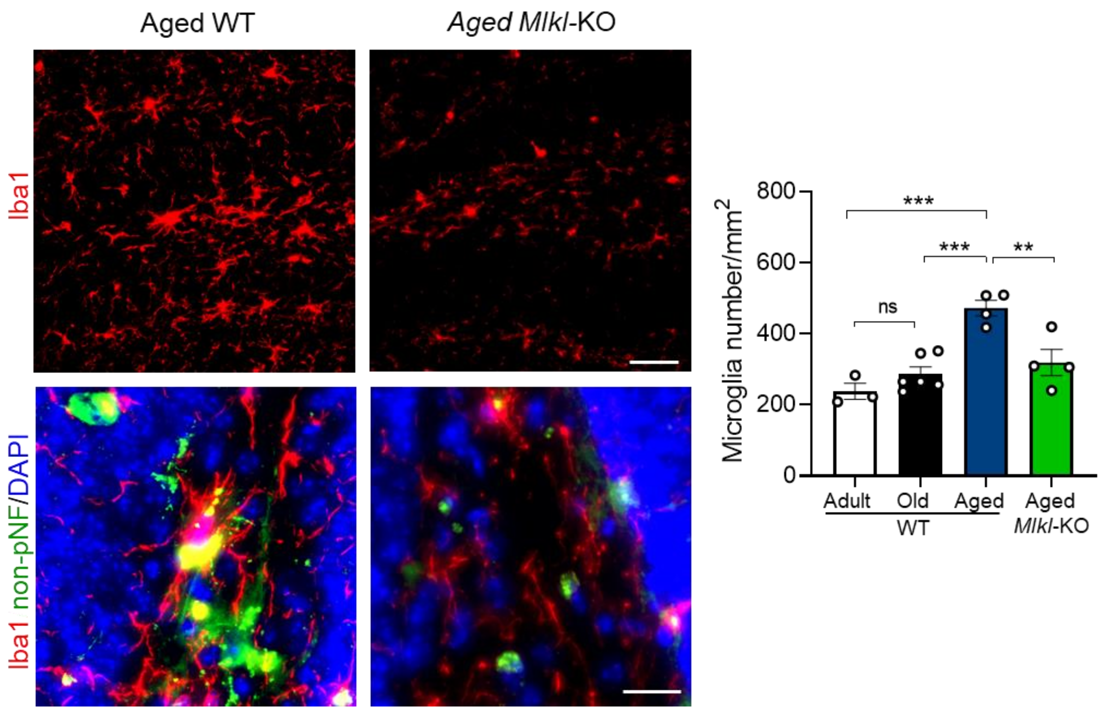


**Fig S5. Microglia activation along aging is prevented by *Mlkl* loss.** Microglia activation was measured by Iba1 immunodetection and quantified as number of positive Iba1 cells in the hilus of the hippocampal DG and normalized to area ($mm^2$), (calibration bar, 50 µm). Microglia overlapping with degenerating axons were identify as Iba1+/non-pNF+ staining (yellow) in the hippocampus of aged mice (calibration bar, 20 µm). Values are the result of the analysis of n=3-5 mice per group. One-way ANOVA with Tukey analysis for multiple comparisons, **p<0.01, ***p<0.005.

Fig. S6

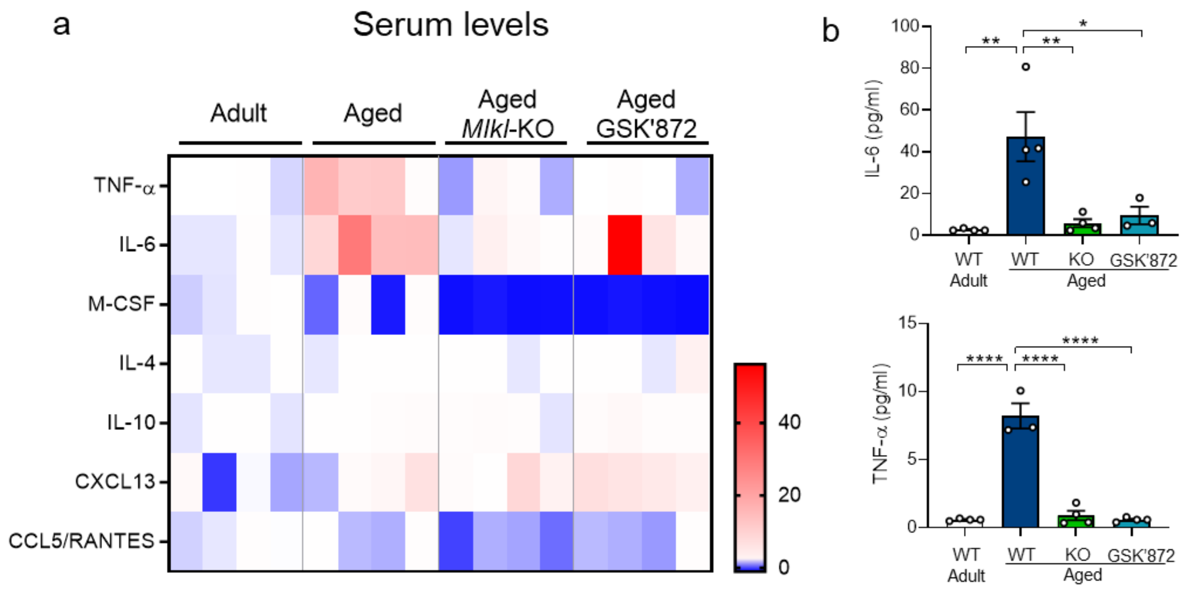


**Fig S6. Systemic decrease of pro-inflammatory cytokines in aged mice with inhibited necroptosis.** Cytokines levels in serum from adult versus aged WT and *Mlkl*-KO mice were assessed by Luminex High Performance Assay. **(a)** The heatmap shows fold change levels of cytokines and chemokines analyzed in the serum of adult versus aged, aged-*Mlkl*-KO and aged-GSK'872 treated mice (n=4, separated in each column). **(b)** Plots represent absolute cytokine levels (pg/ml) for IL-6 and TNF-α in serum samples of different mice groups. Error bars, mean ± SEM, *p:<0.05; **p<0.01; ****p<0.001. Statistical significance was determined by one-way ANOVA with Tukey analysis for multiple comparisons.

Fig. S7

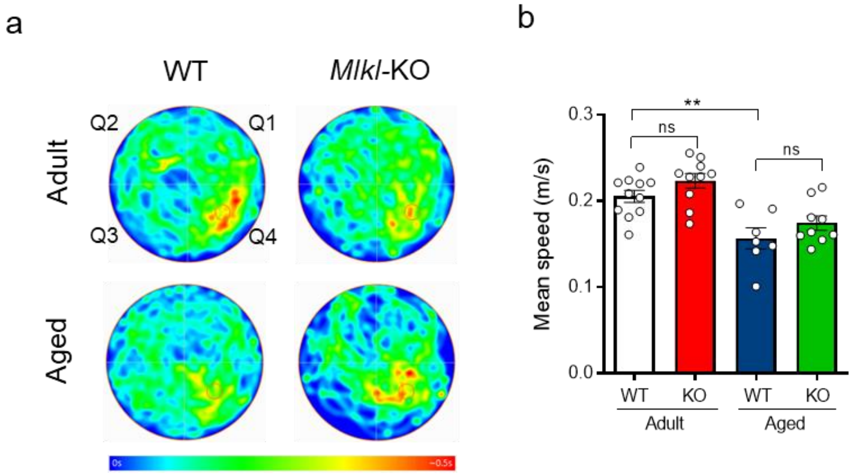


**Fig S7. (a)** Average heatmaps specify in pseudo-color the location of each mice cohort along time during memory testing (day 5). The quadrant Q4 designates the initial location of the hidden platform during training. **(b)** Mean swimming speed of mice during day 5 trial. Values are the result of the analysis of n=7-10 mice per group. One-way ANOVA with Tukey correction for multiple comparison, **p:<0.05.

Fig. S8

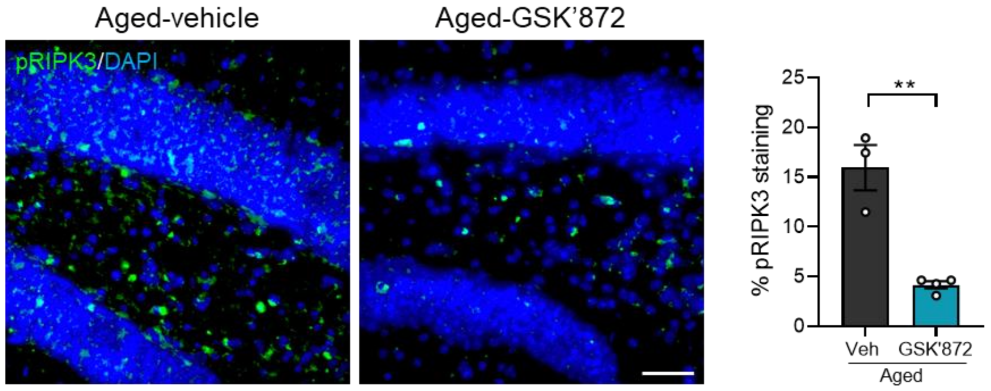


**Fig S8. Decreased pRIPK3 in the hippocampus of aged mice treated with GSK´872.** Efficiency of systemic treatment with the RIPK3 inhibitor and its impact in the brain was evaluated by detecting RIPK3 phosphorylation in the hippocampus of aged mice treated with GSK'872 (calibration bar, 50 µm). RIPK3 activation was measured as the percentage of pRIPK3 stained area. Error bars, mean ± SEM from n=3-4 mice. Statistical significance was determined by T-test with p=0.0018.

Fig. S9

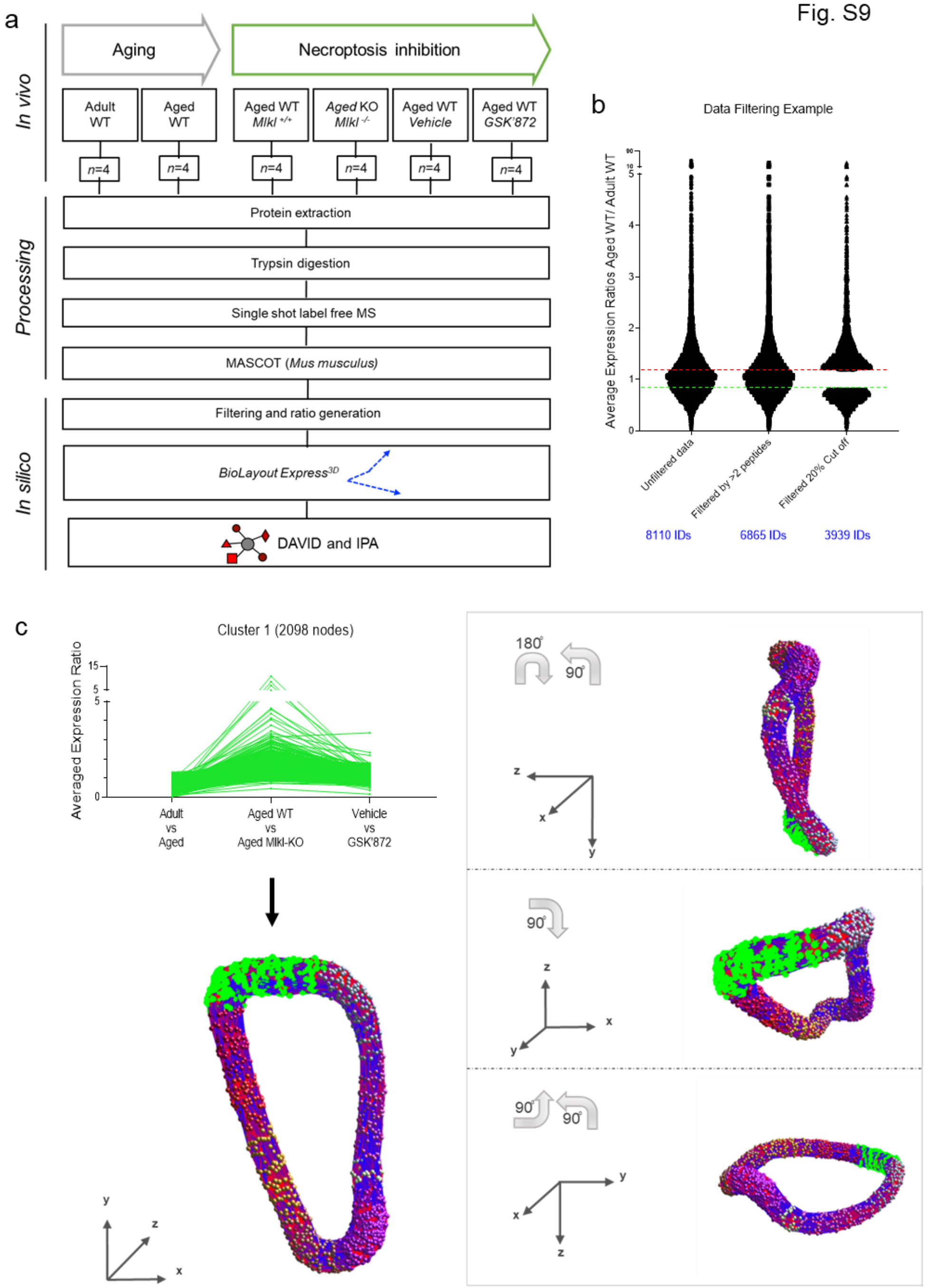


**Fig S9. Proteomic sample processing and data analysis. (a)** Workflow of proteomic analysis and processing of data. **(b)** Data filtering. Proteins identified by ≥ 2 unique peptides and ± 20% of changes were considered for subsequent analysis. **(c)** BioLayout expression analysis allows a 3D representation of the molecular changes between normal aging and necroptosis-targeted aging process. Each sphere represents an individual protein. Clusters (groupings of proteins delineated by color) can be further analyzed with other in silico tools such as DAVID thereby allowing the data to be broken down into more manageable groupings. Cluster 1 is shown in green as clustering example.

Fig. S10

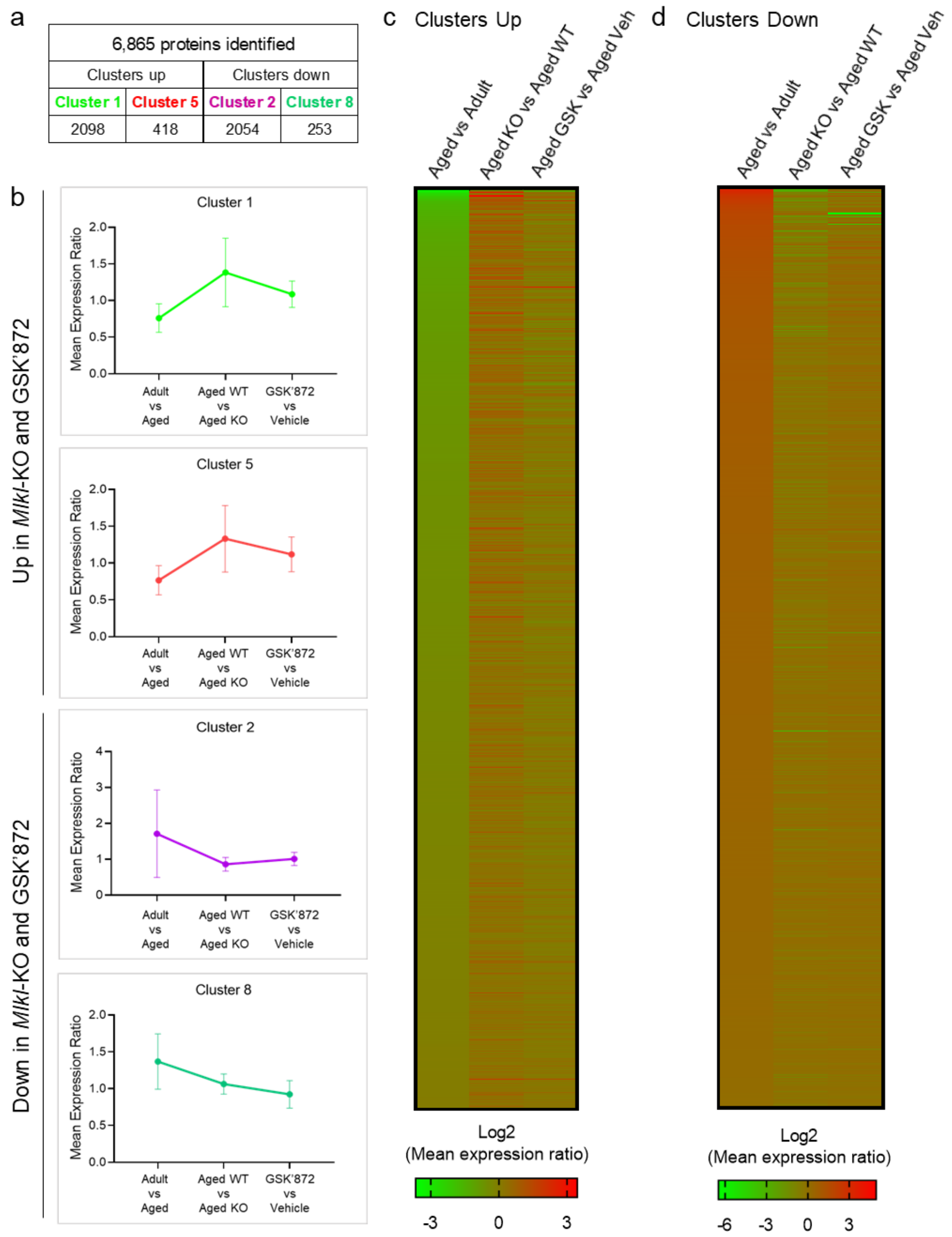


**Fig S10. Proteomic data clustering and analysis. (a)** Table containing information regarding the total number of proteins identified, and those belonging to each cluster of grouping proteins classified as “up” or “down” clusters. **(b)** Classification of clusters “up” and “down” was determined considering the mean expression ratio values and grouped according to their trend to be up- or down-regulated respectively in the genetic (aged *Mlkl*-ko vs aged WT) and pharmacologic (aged-GSK’872 vs aged-vehicle) model of necroptosis inhibition compared with normal aging (adult vs aged). **(c)** Heat maps illustrating the proteomic profile of each cluster show opposing directionality in expression between normal aging and necroptosis-inhibited processes in aged mice. Changes are expressed as log2 of the mean expression ratio.

Fig. S11

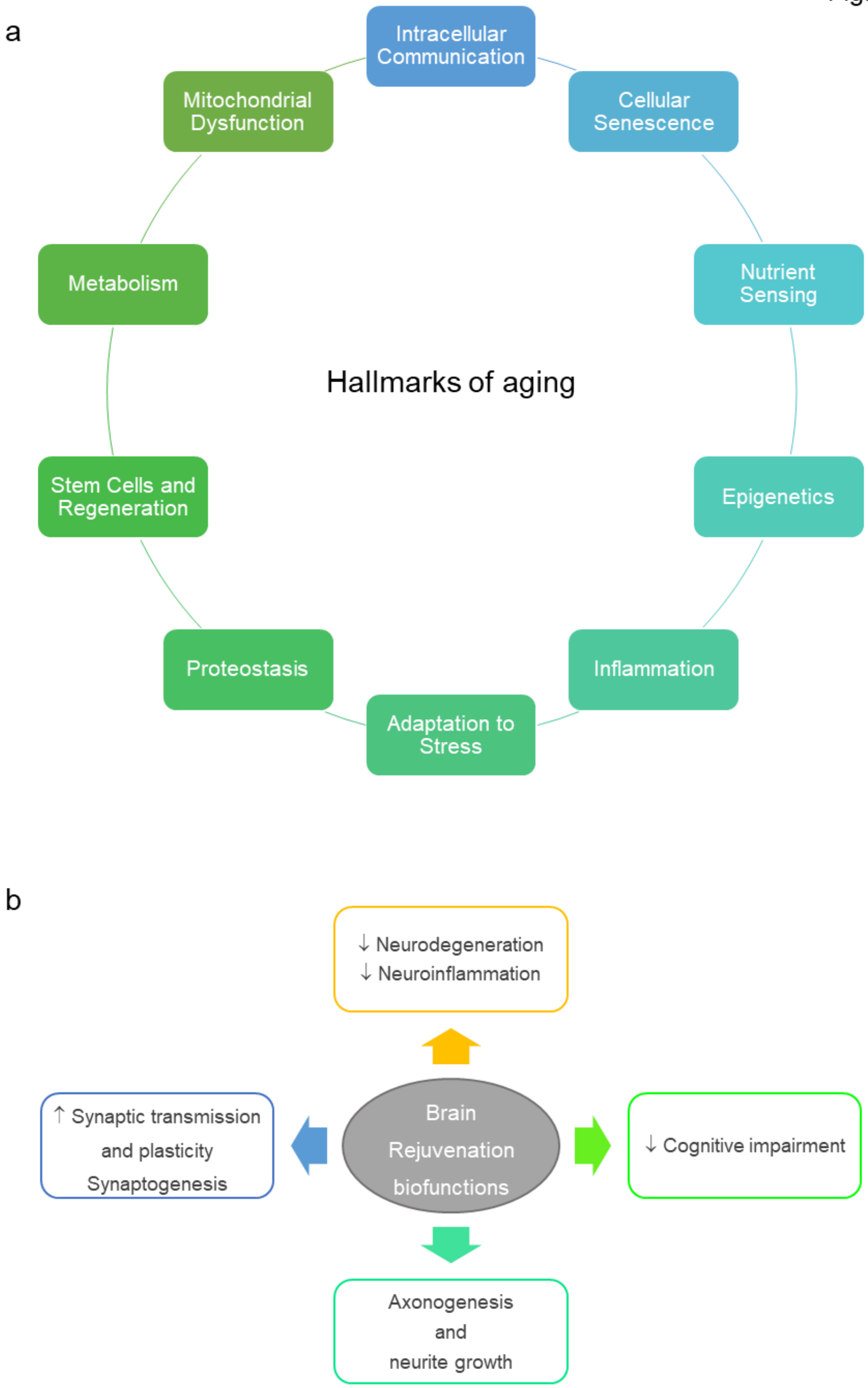


**Fig S11. Pillars of aging and brain rejuvenation hallmarks modulated in the genetic and pharmacologic models of necroptosis inhibition.**

Fig. S12

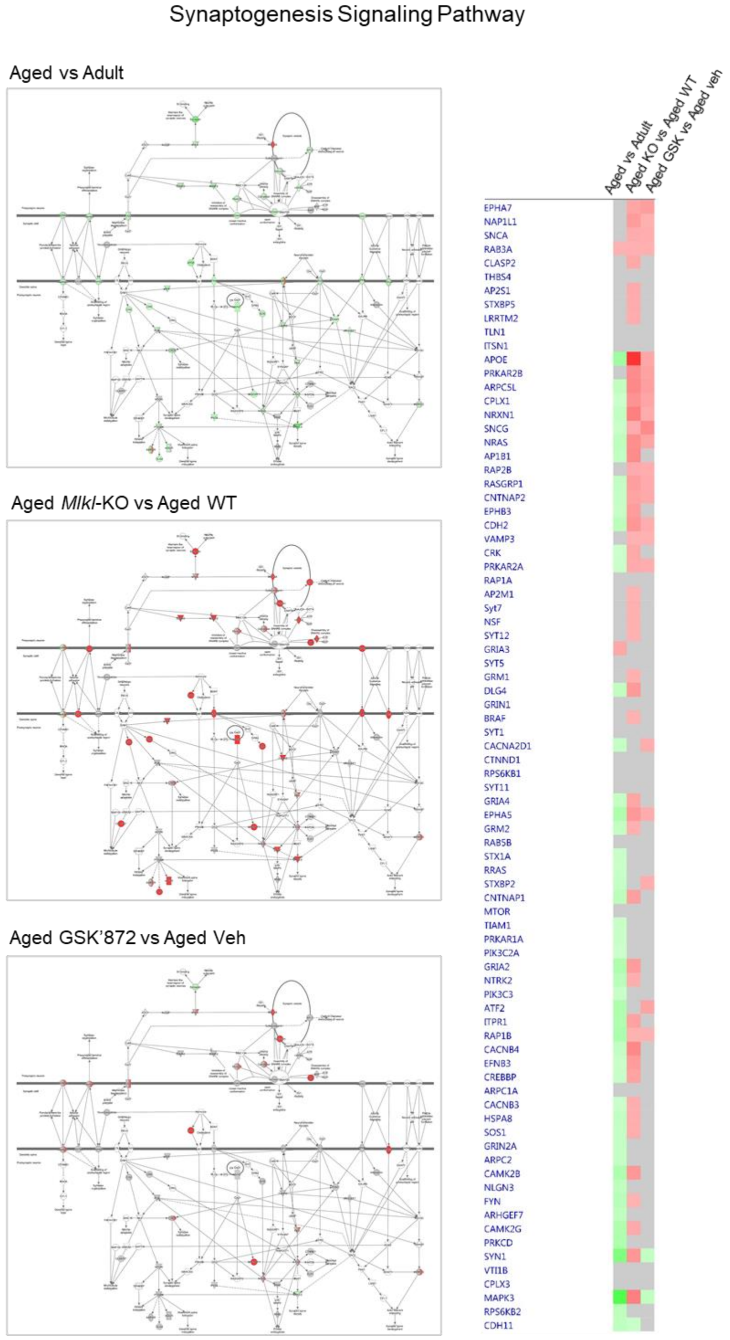


**Fig S12. Schematic illustrating the canonical pathway annotation of "Synaptogenesis signaling pathway". (a)** Top canonical pathway annotations were defined by ranking of absolute z-score between "normal" aging, *Mlkl*-KO and GSK'872 analyses. Intensity of colour represents magnitude of change; red corresponds to upregulation compared to control within each respective analysis, while green represents downregulation. Molecules in grey were identified to be necroptosis-correlative alterations present within input dataset, but fell below the 20% cut-off, while molecules in white were not present within input dataset but changed less than 20% in analysis. Solid connecting lines represent a direct interaction, while dashed connecting lines indicate an indirect interaction. **(b)** Heat map of individual proteins assigned to the canonical pathway "*Synaptogenesis signaling*". Changes between normal aging and *Mlkl*-KO or GSK'872 treated mice were expressed as fold change.

Fig. S13

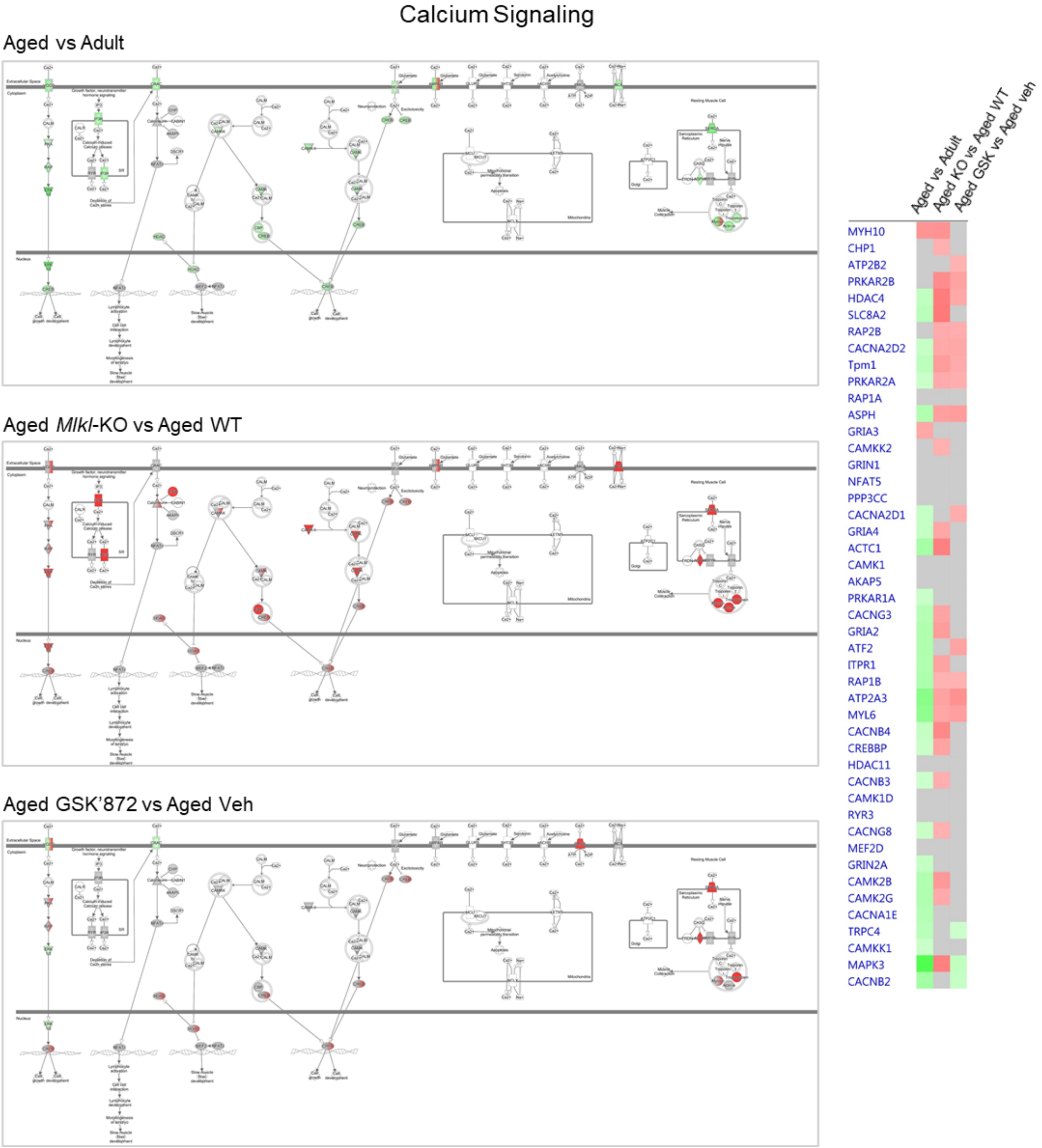


**Fig S13. Schematic illustrating the canonical pathway annotation of “Calcium signaling”. (a)** Top canonical pathway annotations were defined by ranking of absolute z-score between “normal” aging, *Mlkl*-KO and GSK’872 analyses. Intensity of colour represents magnitude of change; red corresponds to upregulation compared to control within each respective analysis, while green represents downregulation. Molecules in grey were identified to be necroptosis-correlative alterations present within input dataset, but fell below the 20% cut-off, while molecules in white were not present within input dataset. Solid connecting lines represent a direct interaction, while dashed connecting lines indicate an indirect interaction. **(b)** Heat map of individual proteins assigned to the canonical pathway “*Calcium signaling*”. Changes between normal aging and *Mlkl*-KO or GSK’872 treated mice were expressed as fold change.

Fig. S14

CREB Signaling in Neurons

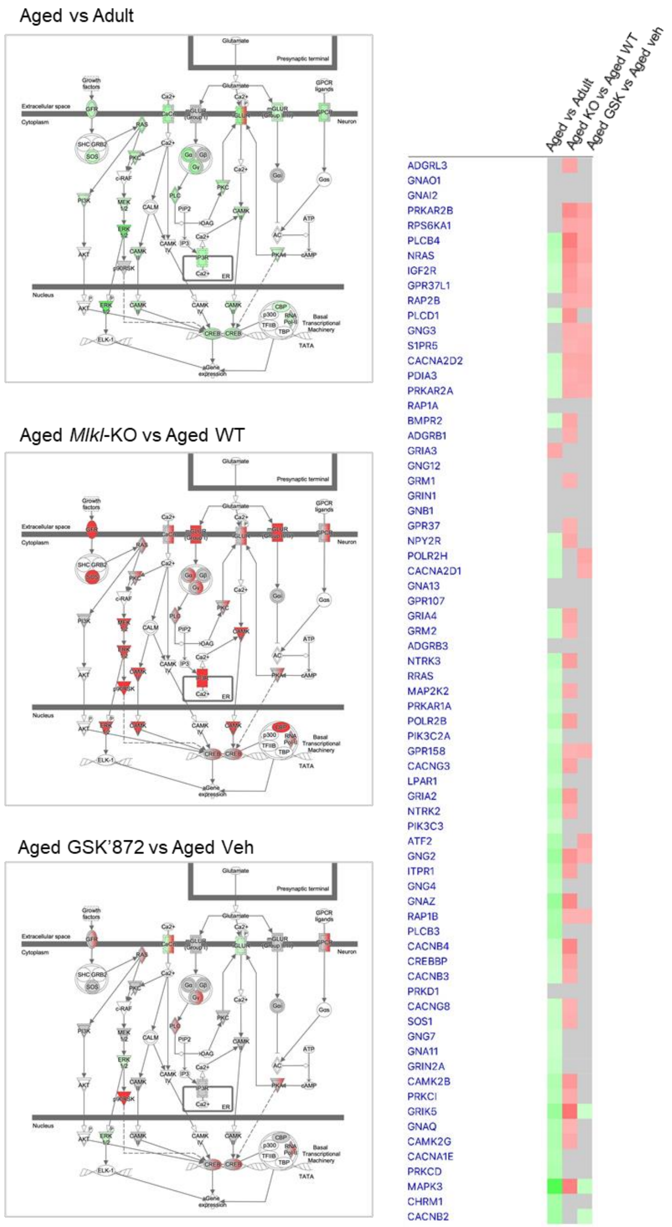


**Fig S14. Schematic illustrating the canonical pathway annotation of "CREB signaling in neurons". (a)** Top canonical pathway annotations were defined by ranking of absolute z-score between "normal" aging, *Mlkl*-KO and GSK'872 analyses. Intensity of colour represents magnitude of change; red corresponds to upregulation compared to control within each respective analysis, while green represents downregulation. Molecules in grey were identified to be necroptosis-correlative alterations present within input dataset, but fell below the 20% cut-off, while molecules in white were not present within input dataset. Solid connecting lines represent a direct interaction, while dashed connecting lines indicate an indirect interaction. **(b)** Heat map of individual proteins assigned to the canonical pathway "*CREB signaling in neurons*". Changes between normal aging and *Mlkl*-KO or GSK'872 treated mice were expressed as fold change.

Fig. S15

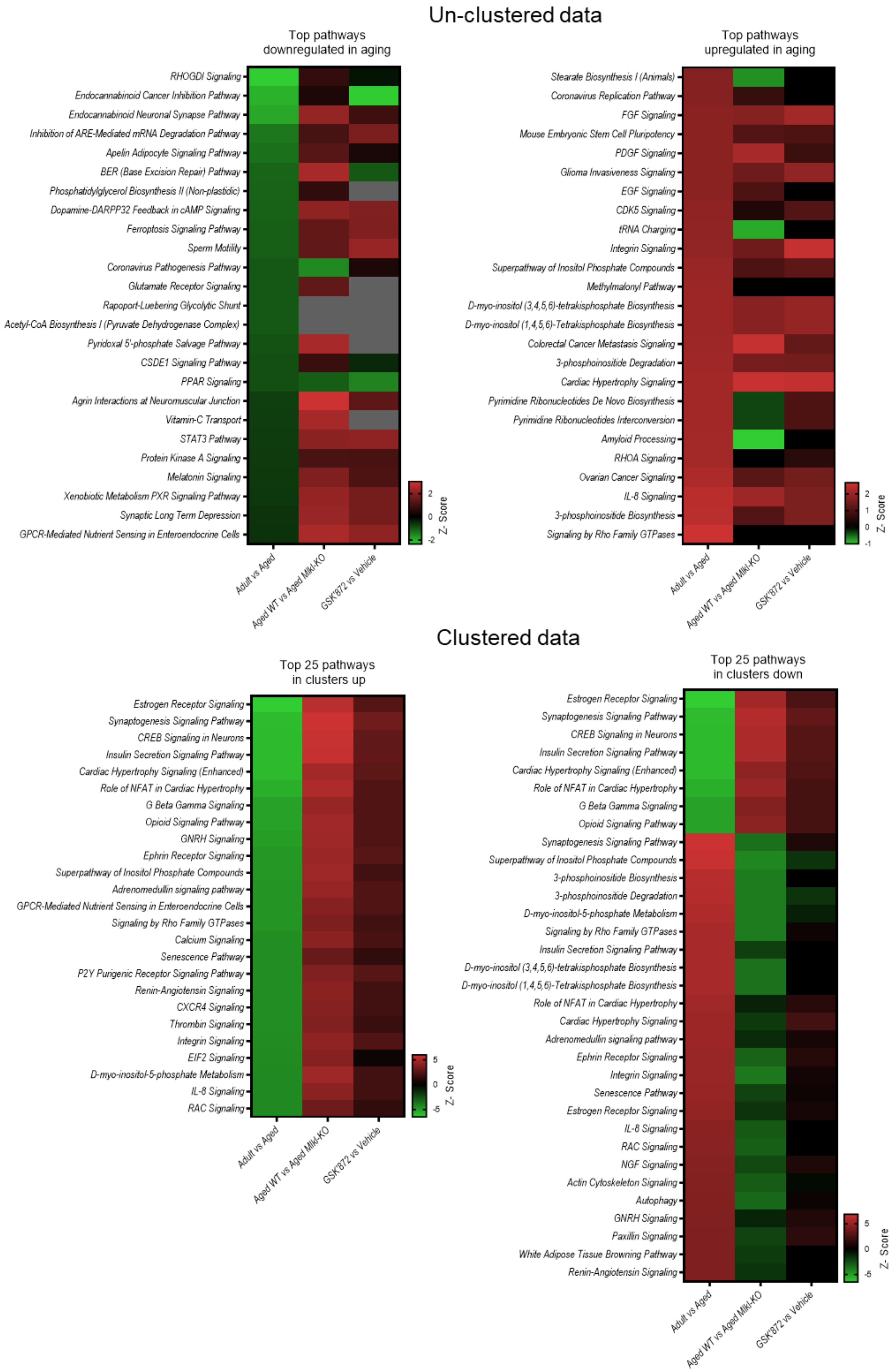
Un-clustered data
Top pathways downregulated in aging
RHOGDI Signaling
Endocannabinoid Cancer Inhibition Pathway
Endocannabinoid Neuronal Synapse Pathway
Inhibition of ARE-Mediated mRNA Degradation Pathway
Apelin Adipocyte Signaling Pathway
BER (Base Excision Repair) Pathway
Phosphatidylglycerol Biosynthesis II (Non-plastidic)
Dopamine-DARPP32 Feedback in cAMP Signaling
Ferroptosis Signaling Pathway
Sperm Motility
Coronavirus Pathogenesis Pathway
Glutamate Receptor Signaling
Rapoport-Luebering Glycolytic Shunt
Acetyl-CoA Biosynthesis I (Pyruvate Dehydrogenase Complex)
Pyridoxal 5'-phosphate Salvage Pathway
CSDE1 Signaling Pathway
PPAR Signaling
Agrin Interactions at Neuromuscular Junction
Vitamin-C Transport
STAT3 Pathway
Protein Kinase A Signaling
Melatonin Signaling
Xenobiotic Metabolism PXR Signaling Pathway
Synaptic Long Term Depression
GPCR-Mediated Nutrient Sensing in Enteroendocrine Cells
Adult vs Aged
Aged WT vs Aged Mlkl-KO
GSK'872 vs Vehicle
2
0
-2
Z- Score
Top pathways upregulated in aging
Stearate Biosynthesis I (Animals)
Coronavirus Replication Pathway
FGF Signaling
Mouse Embryonic Stem Cell Pluripotency
PDGF Signaling
Glioma Invasiveness Signaling
EGF Signaling
CDK5 Signaling
tRNA Charging
Integrin Signaling
Superpathway of Inositol Phosphate Compounds
Methylmalonyl Pathway
D-myo-inositol (3,4,5,6)-tetrakisphosphate Biosynthesis
D-myo-inositol (1,4,5,6)-Tetrakisphosphate Biosynthesis
Colorectal Cancer Metastasis Signaling
3-phosphoinositide Degradation
Cardiac Hypertrophy Signaling
Pyrimidine Ribonucleotides De Novo Biosynthesis
Pyrimidine Ribonucleotides Interconversion
Amyloid Processing
RHOA Signaling
Ovarian Cancer Signaling
IL-8 Signaling
3-phosphoinositide Biosynthesis
Signaling by Rho Family GTPases
Adult vs Aged
Aged WT vs Aged Mlkl-KO
GSK'872 vs Vehicle
2
1
0
-1
Z- Score
Clustered data
Top 25 pathways in clusters up
Estrogen Receptor Signaling
Synaptogenesis Signaling Pathway
CREB Signaling in Neurons
Insulin Secretion Signaling Pathway
Cardiac Hypertrophy Signaling (Enhanced)
Role of NFAT in Cardiac Hypertrophy
G Beta Gamma Signaling
Opioid Signaling Pathway
GNRH Signaling
Ephrin Receptor Signaling
Superpathway of Inositol Phosphate Compounds
Adrenomedullin signaling pathway
GPCR-Mediated Nutrient Sensing in Enteroendocrine Cells
Signaling by Rho Family GTPases
Calcium Signaling
Senescence Pathway
P2Y Purigenic Receptor Signaling Pathway
Renin-Angiotensin Signaling
CXCR4 Signaling
Thrombin Signaling
Integrin Signaling
EIF2 Signaling
D-myo-inositol-5-phosphate Metabolism
IL-8 Signaling
RAC Signaling
Adult vs Aged
Aged WT vs Aged Mlkl-KO
GSK'872 vs Vehicle
5
0
-5
Z- Score
Top 25 pathways in clusters down
Estrogen Receptor Signaling
Synaptogenesis Signaling Pathway
CREB Signaling in Neurons
Insulin Secretion Signaling Pathway
Cardiac Hypertrophy Signaling (Enhanced)
Role of NFAT in Cardiac Hypertrophy
G Beta Gamma Signaling
Opioid Signaling Pathway
Synaptogenesis Signaling Pathway
Superpathway of Inositol Phosphate Compounds
3-phosphoinositide Biosynthesis
3-phosphoinositide Degradation
D-myo-inositol-5-phosphate Metabolism
Signaling by Rho Family GTPases
Insulin Secretion Signaling Pathway
D-myo-inositol (3,4,5,6)-tetrakisphosphate Biosynthesis
D-myo-inositol (1,4,5,6)-Tetrakisphosphate Biosynthesis
Role of NFAT in Cardiac Hypertrophy
Cardiac Hypertrophy Signaling
Adrenomedullin signaling pathway
Ephrin Receptor Signaling
Integrin Signaling
Senescence Pathway
Estrogen Receptor Signaling
IL-8 Signaling
RAC Signaling
NGF Signaling
Actin Cytoskeleton Signaling
Autophagy
GNRH Signaling
Paxillin Signaling
White Adipose Tissue Browning Pathway
Renin-Angiotensin Signaling
Adult vs Aged
Aged WT vs Aged Mlkl-KO
GSK'872 vs Vehicle
5
0
-5
Z- Score

**Fig S15. Top 25 pathways modulated by necroptosis inhibition. (a)** Heat maps illustrating the proteomic profile of the un-clustered data (predicted z-score) of the top pathways up and downregulated in aging. **(b)** Heat maps representing the proteomic profile of the top pathways modulated in the up and down clusters show opposing directionality in expression between normal aging and necroptosis-inhibited processes in aged mice.

Fig. S16

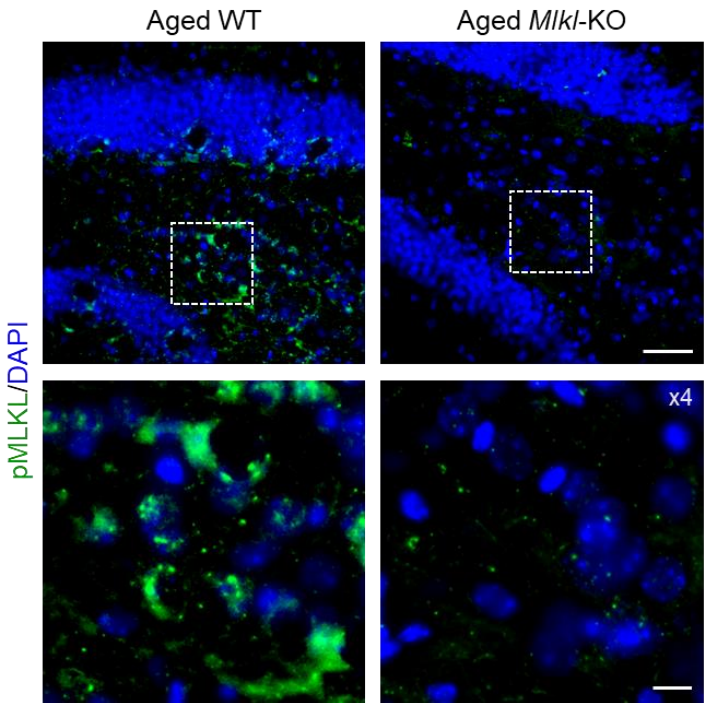


**Fig S16. Phospho-MLKL antibody validation in aged Mlkl-KO mice.** The commercial pMLKL antibody (phospho S345, Abcam, ab196436) was used as a readout of necroptosis activation in hippocampal tissue of aged mice. The pMLKL signal (green) observed in aged mice was validated in the *Mlkl*-KO mouse model by the absence of labeling (calibration bar, 50 µm). White dotted boxes define amplified images at the bottom (x4), showing the staining pattern of pMLKL in the hilus of the hippocampus of aged mice (calibration bar, 10 µm).

**Table S1. Information of analytes evaluated by Luminex High Performance Assay.**

| ANALYTE | BEAD REGION | STANDARD CURVE (pg/ml) | SENSITIVITY (pg/ml) | MAIN FUNCTIONS | REFERENCES |
|---|---|---|---|---|---|
| **CCL2/JE/MCP-1** | 18 | 221 - 53,720 | 134 | Chemokine, regulates migration and infiltration of monocytes/macrophages. | (Barna et al. 1994; Deshmane et al. 2009) |
| **CCL5/RANTES** | 38 | 88.9 - 21,610 | 19.1 | Chemokine, drives mononuclear cells migration across the blood brain barrier (BBB). | (Ubogu et al. 2006; Lanfranco et al. 2018) |
| **CXCL13/BLC/BCA-1** | 21 | 187 - 45,530 | 19.3 | Chemokine, B and T lymphocytes chemoattractant. | (Irani 2016; Pilz et al. 2020) |
| **IFN-gamma** | 33 | 12.3 - 2,990 | 1.85 | Pro-inflammatory cytokine, microglia priming, regulates immune response genes. | (Roy et al. 2020; Monteiro et al. 2015) |
| **IL-1 beta/IL-1F2** | 19 | 221 - 53,780 | 41.8 | Pro-inflammatory cytokine, induces expression of TNF-α and IL-6 to promote neuroinflammation. | (Basu et al. 2004; Shaftel et al. 2008) |
| **IL-2** | 22 | 7.7 - 1,860 | 0.481 | Anti-inflammatory cytokine, regulatory T cells function and survival in autoimmune and inflammatory diseases. | (Ross & Cantrell 2018; Klatzmann & Abbas 2015) |
| **IL-4** | 25 | 56.4 - 13,700 | 53 | Anti-inflammatory cytokine, inhibits IL-1, IL-6 and TNF-α production. Neuroprotective effects. | (Te Velde et al. 1990; Di Benedetto et al. 2017) |
| **IL-6** | 27 | 24.6 - 5,970 | 2.3 | Pro-inflammatory cytokine, induces microglia activation and BBB permeability. Senescence marker. | (Jurk et al. 2012; Erta et al. 2012) |
| **IL-10** | 28 | 11.7 - 2,840 | 8.2 | Anti-inflammatory cytokine, inhibits IL-1, IL-12 and TNF-α production. | (O'Garra & Vieira 2007; Lobo-Silva et al. 2016) |
| **IL-12 p70** | 15 | 42.1 - 10,240 | 12.8 | Pro-inflammatory cytokine required for IFN-γ and TNF-α production. Produces in the brain by microglia. | (Trinchieri 2003; Hofer et al. 2004) |
| **M-CSF** | 45 | 1.85 - 450 | 0.404 | Cytokine, regulates proliferation of macrophages. Microglia polarization towards M1/M2 phenotype. | (Hamilton et al. 2014; Pons & Rivest 2018) |
| **TNF-alpha** | 14 | 3.2 - 790 | 1.47 | The main pro-inflammatory cytokine, activates cell death cascades. Critical for immunoinflammatory response. | (Duque & Descoteaux 2014; Horiuchi et al. 2010) |

**Table S2. Inclusion list.** Mass spectrometry isolation windows for data independent acquisition analysis. Refer to the corresponding method details section for further information.

| Window | Start m/z | End m/z | Width | Centre | Window | Start m/z | End m/z | Width | Centre |
|---|---|---|---|---|---|---|---|---|---|
| **1** | 349.5 | 376.5 | 27 | 363 | **36** | 642.5 | 650.5 | 8 | 646.5 |
| **2** | 375.5 | 396.5 | 21 | 386 | **37** | 649.5 | 657.5 | 8 | 653.5 |
| **3** | 395.5 | 410.5 | 15 | 403 | **38** | 656.5 | 664.5 | 8 | 660.5 |
| **4** | 409.5 | 424.5 | 15 | 417 | **39** | 663.5 | 671.5 | 8 | 667.5 |
| **5** | 423.5 | 435.5 | 12 | 429.5 | **40** | 670.5 | 678.5 | 8 | 674.5 |
| **6** | 434.5 | 445.5 | 11 | 440 | **41** | 677.5 | 685.5 | 8 | 681.5 |
| **7** | 444.5 | 454.5 | 10 | 449.5 | **42** | 684.5 | 692.5 | 8 | 688.5 |
| **8** | 453.5 | 461.5 | 8 | 457.5 | **43** | 691.5 | 700.5 | 9 | 696 |
| **9** | 460.5 | 468.5 | 8 | 464.5 | **44** | 699.5 | 709.5 | 10 | 704.5 |
| **10** | 467.5 | 475.5 | 8 | 471.5 | **45** | 708.5 | 718.5 | 10 | 713.5 |
| **11** | 474.5 | 482.5 | 8 | 478.5 | **46** | 717.5 | 726.5 | 9 | 722 |
| **12** | 481.5 | 489.5 | 8 | 485.5 | **47** | 725.5 | 734.5 | 9 | 730 |
| **13** | 488.5 | 495.5 | 7 | 492 | **48** | 733.5 | 743.5 | 10 | 738.5 |
| **14** | 494.5 | 502.5 | 8 | 498.5 | **49** | 742.5 | 752.5 | 10 | 747.5 |
| **15** | 501.5 | 509.5 | 8 | 505.5 | **50** | 751.5 | 760.5 | 9 | 756 |
| **16** | 508.5 | 516.5 | 8 | 512.5 | **51** | 759.5 | 770.5 | 11 | 765 |
| **17** | 515.5 | 522.5 | 7 | 519 | **52** | 769.5 | 780.5 | 11 | 775 |
| **18** | 521.5 | 529.5 | 8 | 525.5 | **53** | 779.5 | 790.5 | 11 | 785 |
| **19** | 528.5 | 536.5 | 8 | 532.5 | **54** | 789.5 | 801.5 | 12 | 795.5 |
| **20** | 535.5 | 543.5 | 8 | 539.5 | **55** | 800.5 | 812.5 | 12 | 806.5 |
| **21** | 542.5 | 549.5 | 7 | 546 | **56** | 811.5 | 826.5 | 15 | 819 |
| **22** | 548.5 | 555.5 | 7 | 552 | **57** | 825.5 | 839.5 | 14 | 832.5 |
| **23** | 554.5 | 561.5 | 7 | 558 | **58** | 838.5 | 852.5 | 14 | 845.5 |
| **24** | 560.5 | 567.5 | 7 | 564 | **59** | 851.5 | 866.5 | 15 | 859 |
| **25** | 566.5 | 573.5 | 7 | 570 | **60** | 865.5 | 880.5 | 15 | 873 |
| **26** | 572.5 | 580.5 | 8 | 576.5 | **61** | 879.5 | 896.5 | 17 | 888 |
| **27** | 579.5 | 587.5 | 8 | 583.5 | **62** | 895.5 | 916.5 | 21 | 906 |
| **28** | 586.5 | 594.5 | 8 | 590.5 | **63** | 915.5 | 936.5 | 21 | 926 |
| **29** | 593.5 | 602.5 | 9 | 598 | **64** | 935.5 | 958.5 | 23 | 947 |
| **30** | 601.5 | 609.5 | 8 | 605.5 | **65** | 957.5 | 980.5 | 23 | 969 |
| **31** | 608.5 | 615.5 | 7 | 612 | **66** | 979.5 | 1010.5 | 31 | 995 |
| **32** | 614.5 | 622.5 | 8 | 618.5 | **67** | 1009.5 | 1042.5 | 33 | 1026 |
| **33** | 621.5 | 629.5 | 8 | 625.5 | **68** | 1041.5 | 1085.5 | 44 | 1063.5 |
| **34** | 628.5 | 636.5 | 8 | 632.5 | **69** | 1084.5 | 1150.5 | 66 | 1117.5 |
| **35** | 635.5 | 643.5 | 8 | 639.5 | | | | | |
