## Supplementary figures and images for "Necroptosis inhibition counteracts neurodegeneration, memory decline and key hallmarks of aging, promoting brain rejuvenation"

### Graphical Abstract

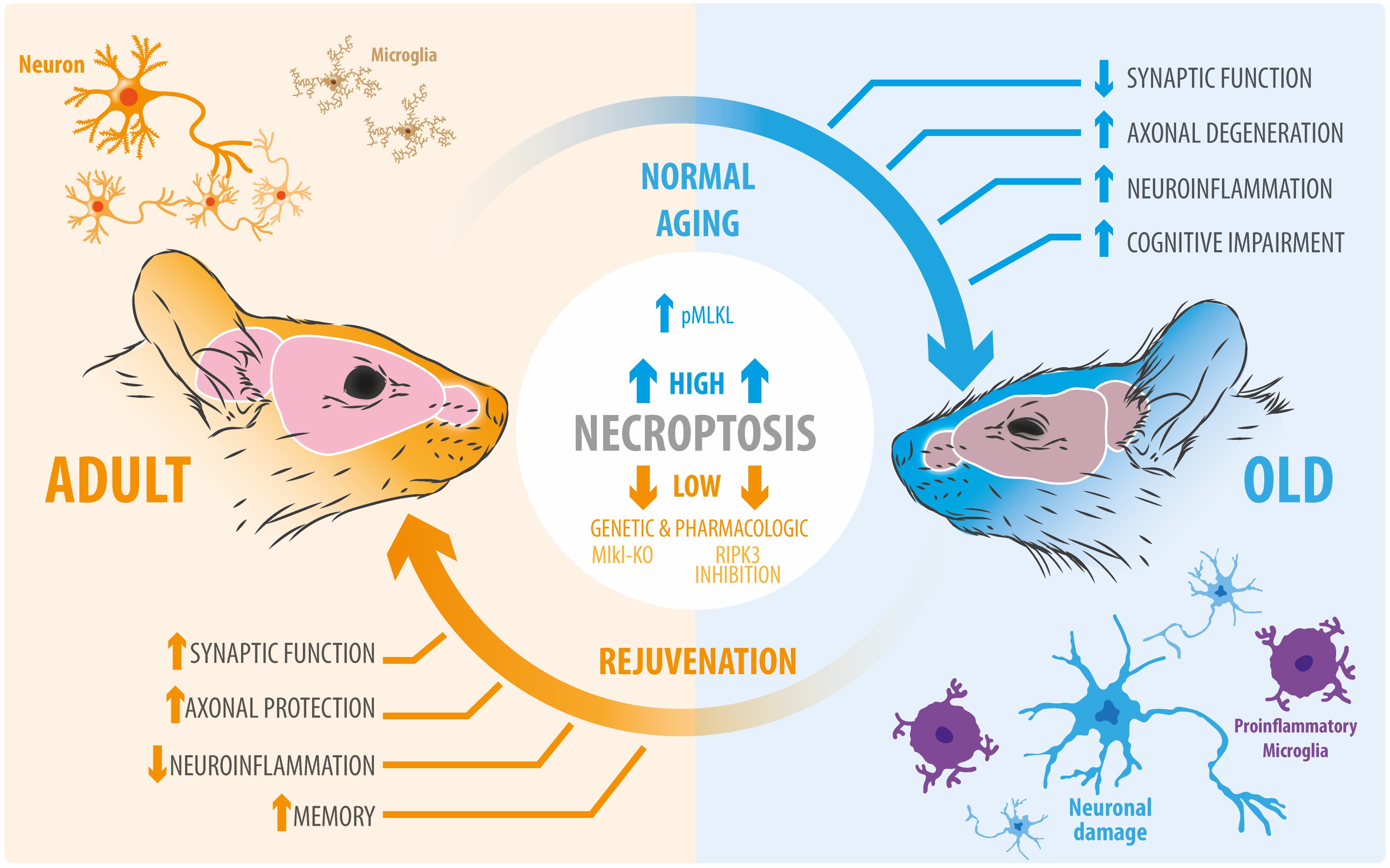
